# Mechanism of phage sensing and abortion by toxin-antitoxin-chaperone systems

**DOI:** 10.1101/2024.02.24.581848

**Authors:** Toomas Mets, Tatsuaki Kurata, Karin Ernits, Marcus J. O. Johansson, Sophie Z. Craig, Gabriel Medina Evora, Jessica A. Buttress, Roni Odai, Kyo Coppieters‘t Wallant, Jose A. Nakamoto, Lena Shyrokova, Artyom A. Egorov, Christopher Ross Doering, Tetiana Brodiazhenko, Michael T. Laub, Tanel Tenson, Henrik Strahl, Chloe Martens, Alexander Harms, Abel Garcia-Pino, Gemma C. Atkinson, Vasili Hauryliuk

**Affiliations:** Department of Experimental Medical Science, Lund University, 221 00 Lund, Sweden; University of Tartu, Institute of Technology, 50411 Tartu, Estonia; Cellular and Molecular Microbiology (CM2), Faculté des Sciences, Université Libre de Bruxelles (ULB), Campus La Plaine, Building BC, Room 1C4203, Boulevard du Triomphe, 1050, Brussels, Belgium; Centre for Bacterial Cell Biology, Biosciences Institute, Newcastle University, Newcastle upon Tyne, NE2 4AX, United Kingdom; Centre for Structural Biology and Bioinformatics, Université Libre de Bruxelles (ULB), Boulevard du Triomphe, Building BC, 1050 Bruxelles, Belgium; Department of Biology, Massachusetts Institute of Technology, Cambridge, MA, USA; Howard Hughes Medical Institute, Massachusetts Institute of Technology, Cambridge, MA, USA; ETH Zurich, Institute of Food, Nutrition and Health, CH-8092 Zürich, Switzerland; Virus Centre, Lund University, Lund, Sweden; Science for Life Laboratory, Lund, Sweden

## Abstract

Toxin-antitoxins (TAs) are prokaryotic two-gene systems comprised of a toxin neutralised by an antitoxin. Toxin-antitoxin-chaperone (TAC) systems additionally include a SecB-like chaperone that stabilises the antitoxin by recognising its chaperone addiction (ChAD) element. TACs have been shown to mediate antiphage defence, but the mechanisms of viral sensing and restriction are unexplored. We identify and characterise two *Escherichia coli* antiphage TAC systems containing HigBA and CmdTA TA units, HigBAC and CmdTAC. The HigBAC is triggered through recognition of the gpV major tail protein of phage λ. Both the ChAD and gpV are recognised by the HigC chaperone through analogous aromatic molecular patterns, explaining the mechanism of activation. We show that the CmdT ADP-ribosyltransferase toxin modifies mRNA to shut down protein synthesis. We establish the modularity of TACs by creating a hybrid broad-spectrum antiphage system combining the CmdTA TA warhead with the HigC chaperone phage sensor.

**Highlights:** *E. coli* HigBAC and CmdTAC are translation-targeting phage immunity TAC systems HigC chaperone recognises phage λ major tail protein to trigger HigBAC toxicity

CmdT ADP-ribosyltransferase toxin abrogates translation through modification of mRNA HigC combined with CmdTA yields hybrid broad-spectrum antiphage defence system

## Introduction

Toxin-antitoxin (TA) systems are diverse two-gene (bicistronic) elements that are ubiquitous in genomes of archaea, bacteria as well as of temperate bacteriophages^1^. While multiple functions have been demonstrated for TAs over the years, their role as abortive infection antiphage defence systems has become a particularly active topic of investigation in recent years^2–4^. Based on the nature of the antitoxin (protein or RNA) and the mechanism of toxin neutralisation (such as formation of inactive toxin-antitoxin complex, protection of the cellular target from the toxin or degradation of the toxin mRNA), TAs are classified into eight groups, from Type I to Type VIII. Type II TAs employ proteinaceous antitoxins that neutralise cognate toxins through the formation of a tight, non-toxic TA complex.

Type II toxins have diverse mechanisms of toxicity, with the most common cellular targets being i) protein synthesis machinery, ii) replication apparatus, iii) cell wall and cell skeleton and iv) nucleotide metabolism^1^. One of the most well-known TA systems is HigBA (<u>H</u>ost inhibition of <u>g</u>rowth), originally identified in *Proteus vulgaris* as a locus that increases the stability of the Rts1 plasmid on which it is encoded^5^. The HigB RNase toxin is a ribosome-dependent mRNA interferase that is neutralised by the N-terminal intrinsically disordered regions of the dimeric HigA antitoxin^6–8^. DarTG is a recently discovered prophage-encoded antiphage defence TA system^9,10^. A member of the ADP-ribosyltransferase (ART) protein family^11^, the DarT toxins link the ADP-ribose moiety of the NAD^+^ cofactor to the amino group of the guanine or thymine base of the DNA^9,12^, which results in inhibition of both RNA and DNA synthesis^10,13^. Detection of the invading phage by DarTG triggers the toxin, thus shutting down cellular transcription and replication in the infected cell to halt virus production^10^.

A variation on the Type II TA theme is provided by tripartite toxin-antitoxin-chaperone (TAC) systems. While TAs are encoded by bicistronic operons, TACs are encoded by tricistronic operons: a ‘classical’ TA gene arrangement followed by a third gene encoding a SecB-like (SecB^TA^) chaperone^14,15^. SecB is a general housekeeping chaperone of Pseudomonadota (Proteobacteria) that predominantly co-translationally assists the folding of diverse proteins, both membrane-targeted and cytoplasmic^16^. SecB^TA^ TAC chaperones are, on the other hand, highly specialised to ensure the stability of otherwise highly labile TAC antitoxins^14^. The strict dependence of the TAC TA units on SecB^TA^-mediated stabilisation is determined by the chaperone addiction (ChAD) element, an intrinsically disordered region (IDR) located at the C-terminus of the antitoxin^17,18^. Rather than mediating antitoxin neutralisation, the ChAD region decreases the antitoxin’s solubility, promoting aggregation and resulting in TA ‘addiction’ to SecB^TA^: the antitoxin and the chaperone act in concert to keep the toxin neutralised. Removal of the ChAD element converts the *Mycobacterium tuberculosis* HigBAC into a SecB^TA^-independent TA system^18^. The interaction between the ChAD region of the *M. tuberculosis* HigA1 antitoxin and SecB^TA^ chaperone HigC is strictly dependent on the aromatic Y114 residue of the ChAD^18,19^. ChAD elements are modular: grafting ChADs on canonical Type II TAs can convert TAs into chaperone-addicted TACs^18^. Finally, the destabilising activity of the ChAD element is conditional on proteolysis of the antitoxin by the ClpXP protease, with *M. tuberculosis* HigBA being rendered non-toxic and SecB^TA^-independent in Δ*clpX* or Δ*clpP* genetic backgrounds^17^.

A recent study by Vassalo and colleagues has established the biological function of two *Escherichia coli* TAC systems: MqsRAC from *E. coli* strain C496_10^18^ and the ART toxin-containing system PD-T4-9 encoded in the hypervariable region of a P2-like phage integrated into the *E. coli* ECOR22 genome^20^. Both of these novel TACs were shown to mediate antiphage defence, and the latter was renamed CmdTAC for <u>C</u>haperone-<u>m</u>ediated <u>d</u>efence <u>TAC</u>^20^. The CmdTAC PD-T4-9 protects *E. coli* from *Tevenvirinae* phage T4, with over-production of the CmdC SecB^TA^ chaperone abrogating protection; the *E. coli* C496_10 MqsRAC protects from *Tevenvirinae* phage T2 but not T4^20^.

In this study we address key outstanding questions in TAC biology: What is the mechanism of toxicity employed by CmdT toxins? How do TAC systems sense phage infection? Using our recent survey of Type II TA diversity^21^ as a starting point, here we identify and validate two novel prophage-encoded TAC systems: HigBAC from *E. coli* strain NT1F31 and CmdTAC from *E. coli* O112ab:H26. We demonstrate HigBAC-mediated defence against siphoviruses (i.e. phages with long noncontractile tails): *Queuovirinae* coliphage Bas25^22^ and *Lambdavirus* λ*vir*. We show that the toxicity of HigBAC is triggered through recognition of the gpV major tail protein of λ*vir* through competition with the ChAD element of the antitoxin HigA. Binding of both HigA and gpV to SecB^TA^ is strictly dependent on specific aromatic residues that compete for the same binding pockets of the chaperone. We uncover the molecular mechanism of anti-*Tevenvirinae* defence by ART toxin CmdTAC: once the TAC system is activated upon phage infection, the CmdT toxin sequence-specifically ADP-ribosylates mRNA to shut down protein synthesis. Finally, by combining the CmdTA toxin-antitoxin warhead with the HigC chaperone phage sensor, we create a hybrid broad-spectrum phage defence system, thus highlighting the evolutionally malleability and modular nature of the TAC architecture. Collectively, our study establishes key principles for phage sensing and restriction by TAC defence systems.

## Results

### *E. coli* NT1F31 HigBAC and *E. coli* O112ab:HH26 CmdTAC are translation-targeting TAC systems that confer phage immunity

In our previous high-throughput analysis of Type II TA systems, the NetFlax algorithm automatically identified TA systems based on their presence in a conserved two-gene architecture^21^. This meant that toxin or antitoxin-like proteins encoded in longer conserved neighbourhoods were not considered. However, when surveying ‘disregarded’ genomic neighbourhoods, we noticed that some toxins from the D2 MqsR-like toxin node of the TA network were associated with an antitoxin followed by a gene encoding a SecB-like protein, suggesting they are TACs, similar to MqsRAC^18,20^. In order to find homologues of these putative TACs that would be suitable for microbiological studies in the *Escherichia coli* host, we searched for relatives of the SecB component in *Enterobacteriacae* and carried out FlaGs gene neighbourhood analysis^23^. This yielded a number of different TAC-like systems in *Enterobacteriacae*, along with *yibN*-*grxC*-*secB*-*gpsA* operons encoding orthologues of the housekeeping *E. coli* SecB^24^ (**Figure S1A**).

We focused on two representatives: HigBAC from *E. coli* strain NT1F31 and a novel tripartite CmdTAC system from *E. coli* O112ab:H26. Both of the systems are located on prophages: a lambdoid phage in the case of *E. coli* NT1F31 HigBAC and an unclassified prophage in the case of *E. coli* O112ab:H26 CmdTAC (**Figure 1A,B**). Importantly, HigBAC is encoded in a variable defence locus downstream of the *cI* repressor^25^ which in the classical λ phage encodes the RexAB phage exclusion system^26^ (**Figure S1B**). Toxicity neutralisation assays establish that both NT1F31 HigBAC and O112ab:H26 CmdTAC are, indeed, *bona fide* TACs. When expressed under the control of an arabinose-inducible PBAD promoter from a pBAD33 plasmid vector, both *E. coli* NT1F31 HigBA (with a weak Shine-Dalgarno sequence) and *E. coli* O112ab:H26 CmdTA (with strong Shine-Dalgarno sequence) toxin-antitoxin units inhibit growth of the BW25113 test strain (**Figure 1A,B**). This demonstrates both the functionality of the toxins and the inability of the full-length antitoxins to neutralise the toxin in the absence of a chaperone. In the case of HigBA, the growth defect caused by TA expression is fully rescued when the cognate HigC chaperone is co-expressed *in trans* from a second plasmid (a pMG25 derivative) under the control of IPTG-inducible PA1/O4/O3 promoter (**Figure 1A**). A partial neutralisation of toxicity is observed for CmdTA co-expressed with CmdC chaperone (**Figure 1B**). Since the operon structure plays an important role in co-translational assembly of protein complexes^27,28^, we tested the toxicity of CmdTAC and HigBAC TAC operons expressed from pBAD33. However, even with the chaperone expressed *in cis* (from the same plasmid), the CmdTA toxicity is not fully suppressed; the likely explanation is excessive toxin expression in our constructs or/and the use of a non-native host strain (**Figure S2**).

**Figure 1.**
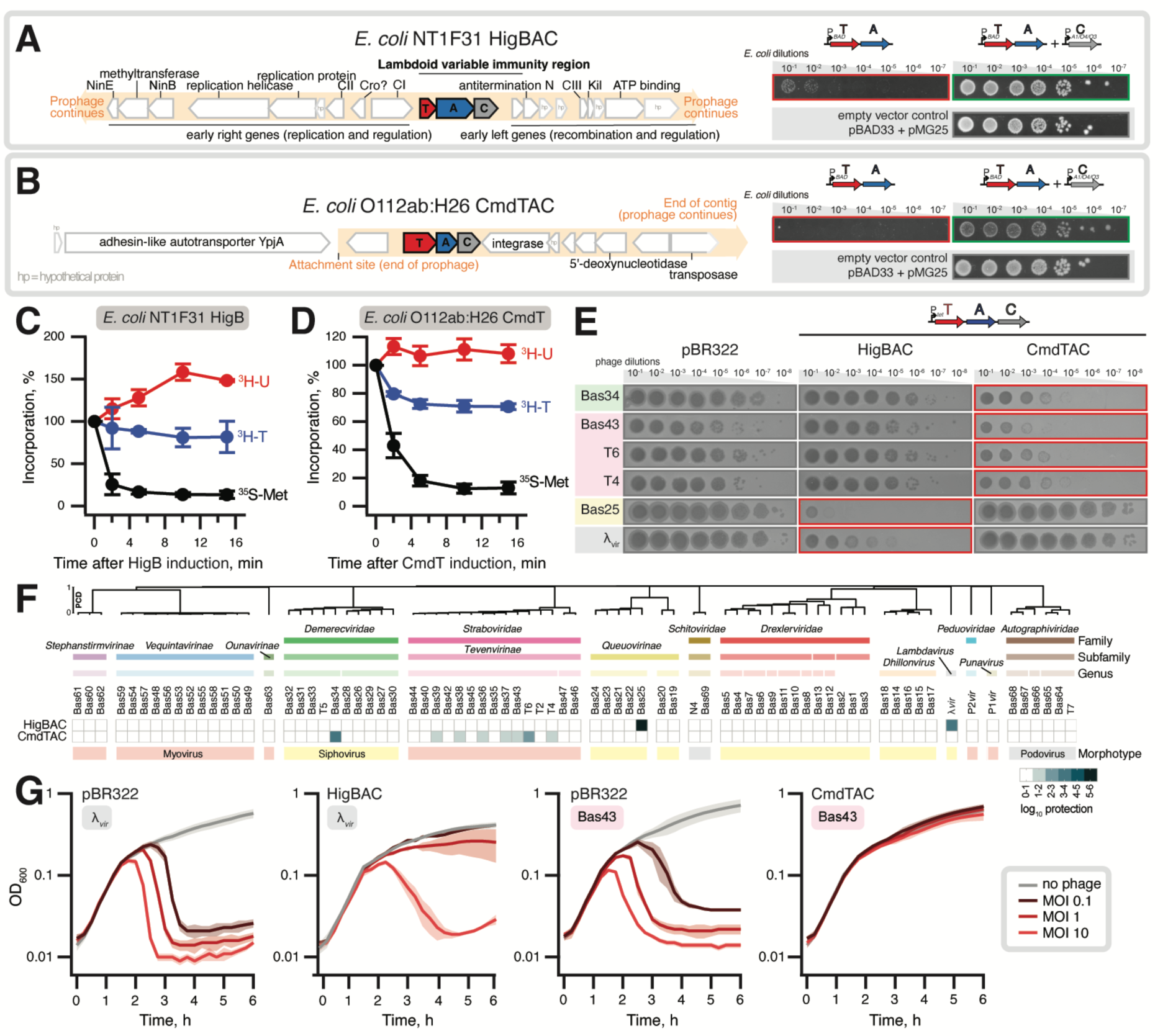
Phage defence by translation-targeting TAC systems: *E. coli* NT1F31 HigBAC and *E. coli* O112ab:HH26 CmdTAC. (**A**,**B**) Gene neighbourhoods of the validated TAC systems: (A) *E. coli* NT1F31 HigBAC and (B) *E. coli* O112ab:HH26 CmdTAC. Both systems are encoded on prophage regions (*left*) and are functional in toxicity neutralisation assays (*right*). To test TAC functionality, *E. coli* BW25113 strains were transformed with empty pBAD33 and pMG25 vectors or their derivatives expressing TAC toxin-antitoxin pairs (expression induced by 0.2% arabinose) and TAC chaperones (induced by 500 µM IPTG), respectively. (**C,D**) Metabolic labelling assays with *E. coli* BW25113 expressing *E. coli* NT1F31 HigB RNase (D) or *E. coli* O112ab:HH26 CmdT ART (C) toxins show specific inhibition of protein synthesis as manifested by a sharp decrease in ^35^S-Met incorporation. (**E,F**) *E. coli* BW25113 cells transformed with either the pBR322 empty vector or either the two pBR322-based plasmids driving the expression of the TAC operons under the control of the constitutive P*tet* promoter were challenged with ten-fold serial dilutions of BASEL^22^ and common laboratory coliphages. (E) Selected phages that were countered by one or other of the TAC systems tested. (F) The results of the full screen shown as a heatmap of log10 protection values. The phage order and dendrogram are defined by hierarchical clustering applied to the Proteome Composition Distance (PCD) matrix (see STAR ★ METHODS section for details). (**G**) Growth of *E. coli* BW25113 carrying the empty vector or the indicated plasmid-encoded TAC system in the presence of λ*vir* or Bas43 phages at MOIs of 0, 0.1, 1 and 10. Additional liquid culture infection experiments with Bas25 and T4 are shown in **Figure S3A-D**.

We used metabolic labelling assays to assess the effects on translation (by following incorporation of ^35^S-methionine in proteins), transcription (incorporation of ^3^H-uridine in RNA) and replication (incorporation of ^3^H-thymidine in DNA) to establish the mechanisms of toxicity employed by the newly identified TACs (**Figure 1C,D**). Given that the Rts1 plasmid-encoded HigB TA toxin is a ribosome-dependent RNase (mRNA interferase)^29^, it is likely that the homologous *E. coli* NT1F31 HigB toxin also degrades mRNA. Indeed, as expected, *E. coli* NT1F31 HigB causes specific and potent inhibition of protein synthesis (**Figure 1C**). The outcome of the labelling experiments with *E. coli* O112ab:H26 ART CmdT toxin was more surprising. In stark contrast to the ART toxin DarT that inhibits replication and transcription^10^, the *E. coli* O112ab:H26 CmdT ART toxin is also a potent and specific inhibitor of protein synthesis, raising the question of the mechanism of CmdT toxicity (**Figure 1D**).

To test the functionality of the identified TACs in antiphage immunity, we performed an antiphage immunity screen using the BASEL coliphage collection as well as a set of commonly used phages^22^ (**Figure 1E,F**). Our collection of double-stranded DNA *Caudoviricetes* viruses included representatives of all the three morphologies: podovirus (short noncontractile tail), siphovirus (long noncontractile tail) and myovirus (long and flexible contractile tail). The tripartite TAC systems were expressed in the BW25113 *E. coli* K12 strain from a pBR322 derivative^30^ under the control of a constitutive Ptet promoter. The *E. coli* NT1F31 HigBAC system provided protection from siphovirus phages such as *Lambdavirus* λ*vir* and *Queuovirinae* Bas25, while *E. coli* O112ab:H26 CmdTAC granted protection against myoviruses from the *Tevenvirinae* subfamily such as Bas43, T4 and T6 as well as a siphovirus from the *Demerecviridae* family (Bas34). Liquid culture infection assays with increasing Multiplicity of Infection (MOI; 0.1, 1 and 10) support the defensive activity of HigBAC against λ*vir* and Bas25 and CmdTAC against Bas43 and T4 (**Figure G** and **Figure S3A-D**). The effects of TAC-mediated defence on phage-mediated culture collapse differ for different experimental systems. Even at MOI of 10, CmdTAC- and HigBAC-expressing *E. coli* strains are virtually immune to Bas43 and Bas25, respectively. At MOI of 10, λ*vir* does eventually cause a collapse of HigBAC-expressing cultures, but more gradually than the rapid collapse in the absence of the defence system. Finally, in the case of CmdTAC-expressing T4-infected *E. coli* cultures, phage-induced lysis is abolished, with the bacterial growth being arrested at the same time points as culture collapse is observed in the cultures of *E. coli* that lack the system. As T4 rapidly degrades bacterial DNA^31^, this possibly reflects efficient abortion of T4’s lytic cycle at the later stages.

Collectively, our results establish these two TAC systems as translation-targeting antiphage immunity systems. Motivated by these results, we set out to answer the following questions: What is the mechanism of CmdT-mediated inhibition of translation? What is the mechanism of phage sensing and restriction by our TAC systems?

### *E. coli* O112ab:H26 ART CmdT ADP-ribosylates mRNA to shut down translation

To validate the direct inhibition of translation by CmdT *in vitro*, we assayed the effect of CmdT on the production of the dihydrofolate reductase (DHFR) reporter protein in a reconstituted *E. coli* cell-free protein synthesis system (PURE)^32^. The toxin was first produced *in situ* in the absence of the NAD^+^ cofactor, and then DHFR production was assayed either in the presence or absence of NAD^+^. The *E. coli* O112ab:H26 ART CmdT toxin potently inhibits DHFR synthesis in a strictly NAD^+^-dependent manner, and, importantly, the 6-biotin-17-NAD^+^ analogue (^biot^NAD^+^) also supports the toxic activity (**Figure 2A**).

**Figure 2.**
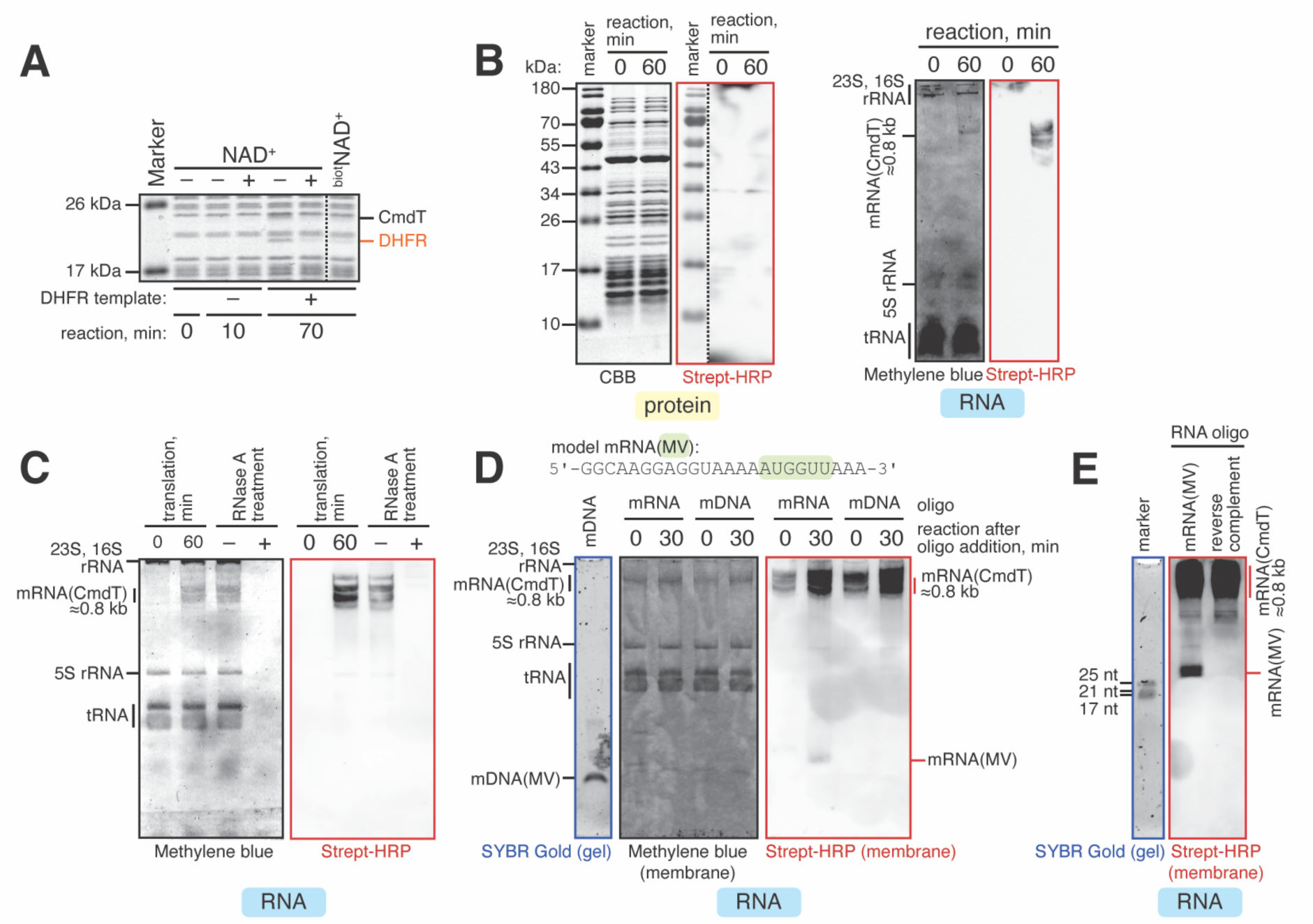
The *E. coli* O112ab:H26 CmdT ART toxin disrupts translation through sequence-specific ADP-ribosylation of mRNA. (**A**) CmdT NAD^+^-dependently abrogates production of DHFR in cell-free expression assays. (**B**,**C**) CmdT modifies the *cmdT*-encoding mRNA in the presence of the biotinylated NAD^+^ (^biot^NAD^+^) substrate. The toxin modifies mRNA(CmdT) but not the protein components of the cell-free expression system (B). The modification signal is sensitive to the addition of RNase A (C). (**D**) CmdT modifies the model mRNA(MV) oligonucleotide but not the corresponding DNA oligonucleotide, mDNA. (**E**) The model RNA(MV) but not its reverse complement RNA is modified by CmdT in the presence of ^biot^NAD^+^.

To identify the specific molecular target modified by CmdT, we used streptavidin-conjugated horseradish peroxidase (streptavidin-HRP) that detects ^biot^NAD^+^. We first produced the CmdT toxin *in situ* in the PURE system in the presence of ^biot^NAD^+^ and after a 60-minute incubation at 37 °C resolved the protein and RNA components on denaturing gels and probed with streptavidin-HRP (**Figure 2B**). While no specific protein signal is detectable, a clear RNA signal is detectable with a size (about 0.8 kb) corresponding to CmdT-encoding mRNA. The signal is sensitive to treatment with single-stranded RNA specific RNase A (**Figure 2C**). Next we used a 24-nucleotide-long model mRNA(MV) (5’- GGCAAGGAGGUAAAAAUGGUUAAA-3’) coding for the MV dipeptide as an RNA substrate. This mRNA is commonly used to assemble ribosomal complexes for biochemical investigations^33^. mRNA(MV) but not the corresponding DNA oligo, mDNA(MV), is readily modified by the toxin (**Figure 2D**). Finally, we used a set of RNA oligos to test whether RNA modification by O112ab:H26 CmdT is sequence-specific. None of the tested homopolymeric RNAs (14-nucleotide-long poly(G), poly(C), poly(U), or poly(A)) are effectively modified, indicating that the toxin recognises a specific sequence motif (**Figure S4**). An RNA oligo that is reverse-complementary to mRNA(MV) was not modified, further suggesting that the modification is sequence-specific (**Figure 2E**). Collectively, our results establish O112ab:H26 CmdT as a sequence-specific mRNA-modifying ART toxin.

### A pattern of conserved aromatic residues of the HigA ChAD determine antitoxin solubility and mediate its recognition by SecB^TA^

Due to more efficient suppression of TA toxicity by SecB^TA^ co-expression in the case *E. coli* NT1F31 HigBAC as compared to *E. coli* O112ab:H26 CmdTAC (**Figure 1B**), the HigBAC is more experimentally amenable for studying the molecular mechanism of ChAD recognition by SecB^TA^. Therefore, we focused on this system. We used AlphaFold-Multimer^34,35^ to co-fold the tetrameric HigC SecB^TA^ in complex with one HigBA TA module (**Figure 3A**, **Figure S5**). At 96 amino acids (residues 122-218), the predicted unstructured HigA ChAD is more than two-fold longer than that of the *M. tuberculosis* HigA1^18^. The ChAD element is predicted to wrap around the SecB^TA^ tetramer, contributing four additional antiparallel β-strands to the conserved β-sheet substructure of the SecB fold. The ChAD β-strands are predicted to occupy the peptide-binding channel of HigC that was previously defined by Xu and colleagues for *Haemophilus influenzae* SecB^36^. Four aromatic residues of the ChAD element – W128, F158, F188 and Y214 – slot into four equivalent pockets of the chaperone subunits involving residues Y39, N41, R49 and D71 of HigC. Analogous pockets accommodating aromatic and hydrophobic ChAD residues were previously observed in the complex of *M. tuberculosis* SecB^TA^ bound to a short ChAD peptide fragment^19^.

**Figure 3.**
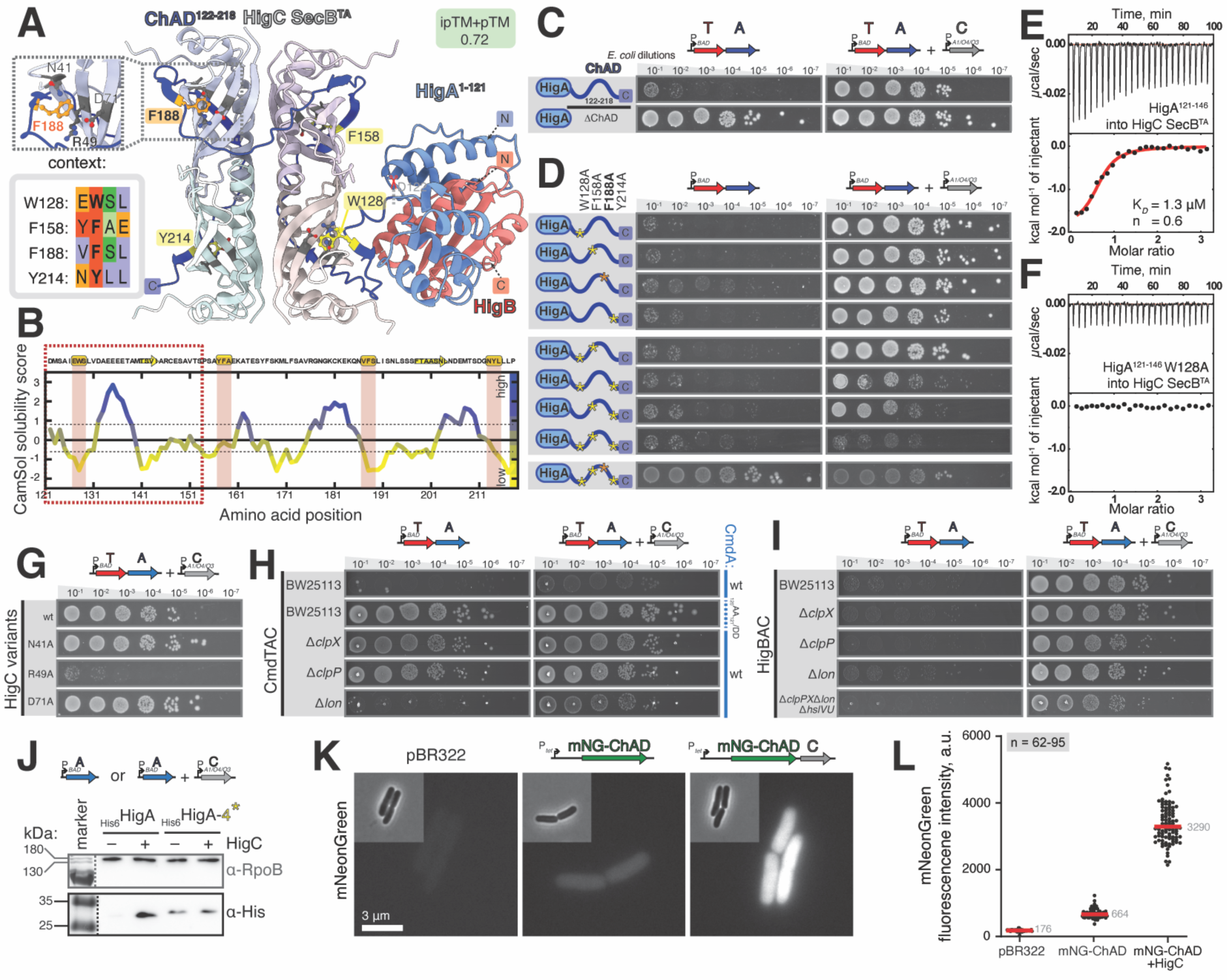
The aromatic residues of the ChAD element play key roles in HigC-dependent neutralisation of the HigB toxin by HigA. (A) An AlphaFold model of HigBA TA unit in complex with the HigC tetramer. The four aromatic residues of the HigA ChAD element that are predicted to interact with the four equivalent pockets of the HigC4 tetramer are highlighted. **(B)** CamSol solubility analysis of the HigA ChAD region, residues 122-218. The four putative SLiM motifs are highlighted while the ChAD section used for ITC experiments is marked with a dashed box. (**C**,**D**) Toxicity assays with wild-type and ChAD-mutated HigA antitoxin variants. Removal of the ChAD element renders the HigBA TA unit non-toxic, indicative of efficient HigA stabilisation (C). Effects of individual substitutions of the four aromatic residues of the ChAD element on HigBA toxicity (D, *left*) and its suppression through co-expression of the HigC (D, *right*). HigBA and HigC expression was induced by 0.2% arabinose and 50 µM IPTG, respectively. (**E**,**F**) Binding of the wild-type (E) and W128A-substituted (F) HigA^121–146^ peptide to HigC as monitored by ITC. The stoichiometry is calculated per HigC protein chain. The test peptide contains two aromatic residues (W128 and F158) out of four found in the full ChAD element. (**G**) Suppression of HigBA toxicity through IPTG-induced co-expression of HigC variants carrying amino acid substitutions in the pocket that is predicted to recognise the aromatic residues of the HigA ChAD. The R49A HigC variant is unable to suppress the HigBA toxicity. (**H,I**) TA and TAC toxicity assays in *E. coli* BW25113 protease-deficient strains. Indicative of antitoxin degradation by ClpXP, the CmdTA TA unit is non-toxic in Δ*clpP* and Δ*clpX* but not Δ*lon* strains (H, *left*). The CmdTA TA toxicity is abrogated upon the Asp-Asp substitution of the C-terminal Ala-Ala motif of CmdA that is recognised by ClpX (H). The HigBA TA is toxic in all of the tested backgrounds, suggesting that proteolytic degradation is not essential for ChAD-mediated destabilisation of the antitoxin (I, *left*). TA and chaperone expression was induced by 0.2% arabinose and 500 µM IPTG, respectively. (**J**) Immunoblotting analysis of *E. coli* BW25113 cells expressing either wild-type or substituted (4*) N-terminally 6His-tagged HigA, either in the absence or in the presence of HigC. (**K**,**L**) Fluorescence and phase contrast microscopy of *E. coli* BW25113 cells transformed with an empty pBR322 vector or pBR322 derivatives expressing either mNeonGreen-ChAD alone or mNeonGreen-ChAD together with HigC (K). Quantification of mNeonGreen fluorescence for individual cells from the same imaging dataset (n = 62 to 95 cells) (L).

The aromatic residues of HigA ChAD are conserved, supporting their functional importance (**Figure S6A**). Intrinsically disordered regions (IDRs) often recognise their target proteins through recognition of Short Linear Motifs (SLiMs) or/and shorter specific sequence features, such as clusters of aromatic of charged residues^37^. Closer inspection of the four aromatic residues reveals that i) there is a loose SLiM consensus associated with the residues: with the exception of F188, aromatic residues are preceded by polar residue and followed by an amino acid with a small side-chain, ii) the three potential SLiMs are regularly interspaced across the ChAD with a step of ≍30 amino acids, and iii) as predicted by CamSol^38^, F188 is located in a region with a high propensity for aggregation (**Figure 3B**) thus suggesting a key role in the destabilising activity of the ChAD element.

Removal of the predicted ChAD region (Δ122-218) renders the *E. coli* NT1F31 HigA antitoxin able to efficiently neutralise the HigB toxin in the absence of the chaperone, behaving like a classical *higBA* TA pair (**Figure 3C**). To validate our AlphaFold predictions of specific interactions, we next targeted the four aromatic residues of the HigA ChAD element. First, we substituted each individually for alanine and tested the toxicity of the resultant mutant *higBA* TA units in the presence or absence of the chaperone (**Figure 3D**). The only mutant *higBA* TA variant that had a phenotype was the F188A substitution that is predicted to be a part of the CamSol-predicted aggregation-prone patch. This TA variant is slightly less toxic than the wild-type (**Figure 3D**, *left*), while the HigBA toxicity is still efficiently neutralised by HigC (**Figure 3D**, *right*). Next, we substituted W128, F158 and Y214 in combinations, leaving out F188. While none of the tested pairwise substitutions abolished the HigC-mediated suppression of toxicity, substitution of the three residues for alanine completely abrogated the effect of HigC co-expression (**Figure 3D**, *right*). This suggests that while they are not individually essential for SecB^TA^ addiction, these residues collectively participate in the interaction through avidity effects. Finally, the tetra-substituted variant has a dramatically reduced toxicity which does not need to be suppressed by HigC. This suggests that the four residues are collectively essential for rendering the ChAD-tagged antitoxin aggregation-prone and chaperone-addicted. Next, we used isothermal titration calorimetry (ITC) to test the HigC-mediated recognition of a peptide mimicking the first of the four repetitive SLiM elements of the ChAD region (**Figure 3E**,**F**, **Table 1**). The peptide corresponding to residues 121-148 of HigA binds HigC with low-μM affinity (K*D* = 1.3 μM, n = 0.6; n is calculated per HigC monomer). The W128A substitution targeting the aromatic core of the SLiM abrogates the interaction, directly validating specificity of the interaction.

**Table 1.** Binding parameters of gpV and HigA fragments to HigC. . Experiments were performed with either 30 µM HigC (titrated with 350 µM gpV^1-160^) or 16 µM HigC (titrated with 160 µM gpV^48–78^ or HigA^121-146^, either wild-type or substituted variants). In the competition experiment 12 µM HigC supplemented with 18 µM HigA^121-146^ was titrated with 160 µM gpV^48-78^ similarly supplemented with 18 µM HigA^121-146^ in order to prevent the dilution of the ChAD-mimicking peptide. The binding parameters were determined by fitting the ITC data to a single interaction model. Data represent mean values ± s.d., N.D. stands for ‘not detectable’. The presented titrations are background-subtracted.

| Titrations | $K_D$ ($\mu\text{M}$) | $\Delta H$ (kcal/mol) | $-T\Delta S$ (kcal/mol) | $\Delta G$ (kcal/mol) | Molar Ratio (n) |
| --- | --- | --- | --- | --- | --- |
| gpV<sup>1-160</sup> into HigC | N.D. | N.D. | N.D. | N.D. | N.D. |
| gpV<sup>48-78</sup> into HigC | 0.25 | -3.8 | -5.3 | -9.1 | 0.47 |
| gpV<sup>48-78</sup><sub>W67A</sub> into HigC | N.D. | N.D. | N.D. | N.D. | N.D. |
| gpV<sup>48-78</sup><sub>61DDED64/AAAA</sub> into HigC | 7.0 | -4.5 | -2.6 | -7.1 | 0.49 |
| HigA<sup>121-146</sup> into HigC | 1.3 | -1.9 | -6.3 | -8.2 | 0.6 |
| HigA<sup>121-146</sup><sub>W128A</sub> into HigC | N.D. | N.D. | N.D. | N.D. | N.D. |
| (gpV<sup>48-78</sup>+HigA<sup>121-146</sup>) into (HigC+HigA<sup>121-146</sup>) | 6.7 | -5.6 | -1.6 | -7.2 | 0.5 |

Finally, we probed the pockets of the HigC chaperone that are predicted to recognise the aromatic residues of the ChAD element. While alanine substitution of N41 and D71 do not affect the HigC-mediated neutralisation of HigBA, the R49A variant is unable to rescue the growth defect of HigBA-expressing cells (**Figure 3G**).

**Inactivation of the *E. coli* NT1F31 HigA antitoxin does not rely on the ClpXP protease** The protease ClpXP recognises specific motifs, or degrons, that mark a protein for degradation. The most well-characterised ClpXP degron is the SsrA-tag. The key element of this tag is the C-terminal Ala-Ala-COO^-^ moiety^39,40^. The ChAD of *M. tuberculosis* HigA1 terminates with the similarly aliphatic Val-Ala dipeptide degron, marking the antitoxin for ClpXP-mediated degradation^17^. Substitution of the C-terminal Val-Ala motif of *M. tuberculosis* HigA1 for Asp-Asp abolishes antitoxin degradation by ClpXP, rendering the well-neutralised HigBA1 toxin-antitoxin unit non-toxic^17^.

The ChAD element of CmdA terminates with Ala-Ala, suggesting targeting by ClpXP. The *E. coli* NT1F31 HigA on the other hand terminates with Leu-Pro, which makes it an unlikely substrate for ClpXP. To test the role of ClpXP in the function of the two TAC systems, we substituted the terminal AA motif of CmdA for DD and tested the toxicity of the resultant CmdTA construct in wild-type BW25113 *E. coli* (**Figure 3H**). Consistent with ClpXP being responsible for the instability of wild-type CmdA for degradation, the 120AA121/DD mutant CmdTA variant is nontoxic. Next, we tested the toxicity of the two TA modules in BW25113. *E. coli* Δ*clpP*, Δ*clpX* and Δ*lon* strains; the latter strain was used as a specificity control. As expected, the CmdTA unit is not toxic in Δ*clpP* and Δ*clpX* – but is toxic in Δ*lon* – *E. coli* strains, supporting the direct role of ClpXP in CmdTAC triggering through antitoxin degradation (**Figure 3H**). As expected from its likely lack of a degron in the ChAD element, HigAB is toxic in all tested protease-deficient backgrounds, including Δ*clpPX* Δ*lon* Δ*hsIVU* (**Figure 3I**). This suggests that HigBAC activation does not rely on the proteolytic activity of ClpX, Lon or HslVU. Rather, it could rely on antitoxin aggregation or/and clearance by an as yet undetermined protease(s).

To directly assay the effects of HigC on HigA abundance and solubility, we used immunoblotting (**Figure 3J**) and fluorescence microscopy (**Figure 3K,L**). First, we tested the effects of HigC co-expression as well as simultaneous alanine substitutions of the four aromatic residues of the HigA ChAD element (W128A, F158A, Y214A and F188A; referred to as the 4*) on the expression levels of N-terminally His6-tagged HigA (**Figure 3J**). In the absence of HigC, wild-type HigA is not detectable by α-His6 immunoblotting; co-expression of HigC results in a strong and specific α-His6-signal. Conversely, the 4* ChAD-mutated HigA is stable regardless the presence or absence of HigC. These results demonstrate that HigC:ChAD interaction is essential stabilising otherwise highly liable HigA and, second, that the aromatic residues of the ChAD element are crucial for its degron function. Next, we imaged *E. coli* cells expressing the C-terminally ChAD-tagged monomeric yellow-green fluorescent protein mNeonGreen^41^ with or without co-expression of HigC; *E. coli* transformed with empty pBR322 vector were used as a control (**Figure 3K,L**). In good agreement with the immunoblotting experiments, HigC co-expression increased the fluorescent signal 5-fold. In good agreement with immunoblotting experiments, HigC co-expression increased the fluorescent signal 5-fold. In the absence of HigC, the mNeonGreen-ChAD fluorescent signal is evenly distributed though the cell with no aggregate formation detected, indicating that the association of HigA protects the antitoxin from proteolytic destruction rather than aggregation.

### Toxic activity of the *E. coli* NT1F31 HigBAC system is trigged by direct sensing of the λ phage major tail protein gpV

To discover the nature of the phage triggers that activate the *E. coli* NT1F31 HigBAC defence system, we isolated spontaneous escape mutants of the λ*vir* phage that can overcome HigBAC- mediated immunity. We sequenced four λ*vir^escape^* mutant phages immune to HigBAC, and all shared the same S54P substitution in the major tail protein gpV (**Dataset S1**). In the absence of HigBAC, wild-type λ*vir* and λ*vir^escaspe_1^* are similarly active against *E. coli*; in the presence of HigBAC only the wild-type λ*vir* is restricted (**Figure 4A, Figure S3E,F**). Given that structural proteins are well-established triggers of antiphage defence systems^42,43^, we have focused our attention of the gpV as a candidate HigBAC trigger. S54 is located in a highly flexible region of the β2-β3 loop (residues 50-78) of the N-terminal domain of gpV (gpVN, residues 1-160) that mediates polymerization of the phage tail^44,45^. Thus, the S54P substitution would likely not significantly alter the structural properties of gpV given the intrinsic flexibility of the region.

**Figure 4.**
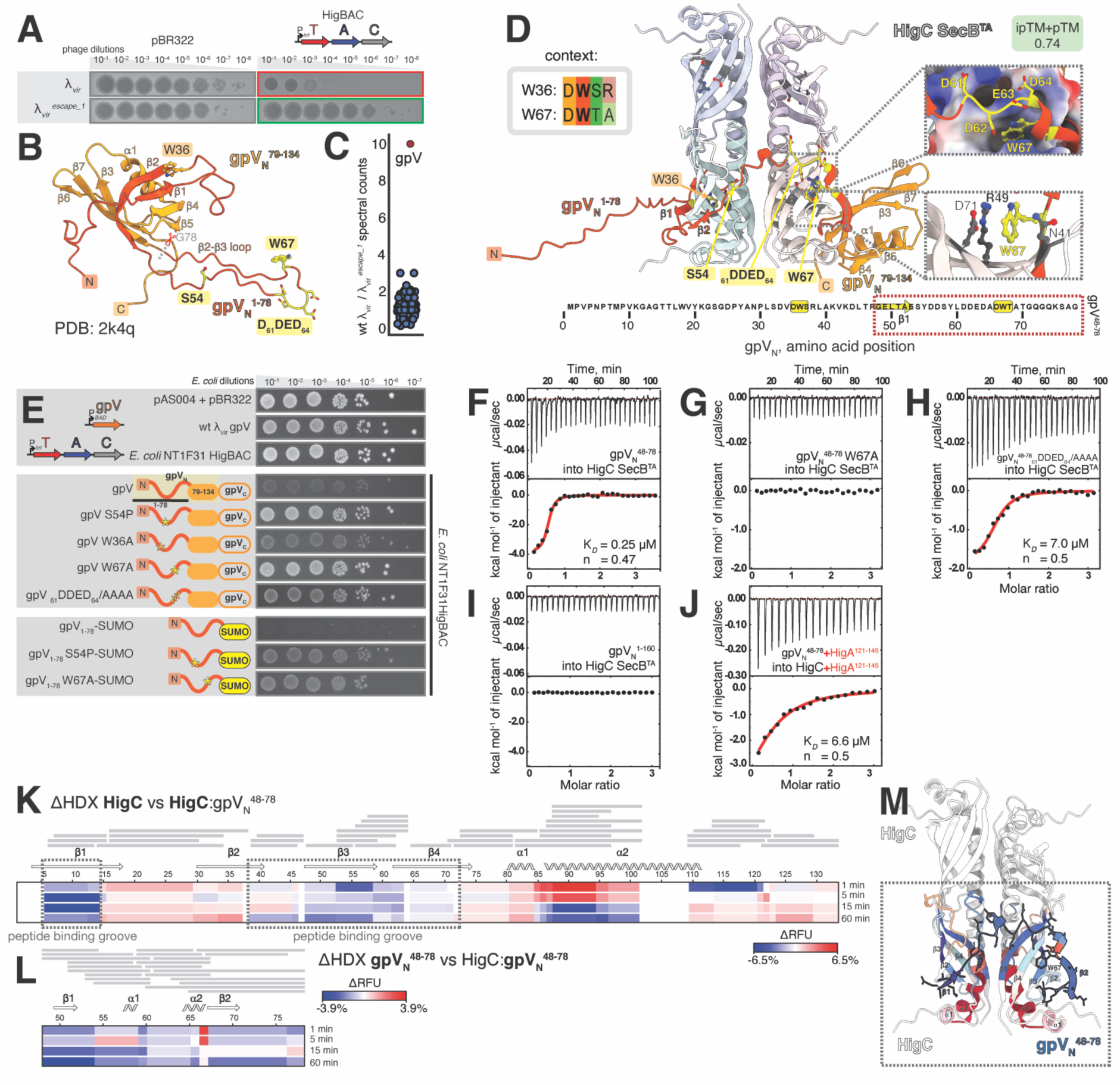
The λ phage major tail protein gpV triggers the E. coli NT1F31 HigBAC system. (A) Constitutive Ptet-driven expression of HigBAC provides protection against λ*vir* but not the λ*vir^escape_1^* escape mutant carrying an *S54P* substitution in *gpV*. Complementary liquid culture infection experiments are shown on **Figure S3E**,**F**. **(B)**Experimentally determined structure of gpVN (PDB 2k4q)^45^. **(C)** Comparative IP-MS/MS analysis of HigBA_FLAG-HigC pulldown samples purified from *E. coli* culture infected with λ*vir* and λ*vir^escape_1^* mutant reveals relative enrichment of gpV in the wild-type sample. **(D)** An AlphaFold model of the λ gpVN domain in complex with the HigC tetramer. The gpVN^48–78^ section used for ITC experiments is highlighted with a dashed box. **(E)** Induction of HigBAC toxicity though co-expression of wild-type and engineered gpV variants. While expression of either HigBAC or gpV alone has no effect, their co-expression results in synthetic toxicity. Co-expression of HigBAC with the escape S54P mutant variant of gpV fails to trigger TAC-mediated toxicity. No triggering is observed for W67A-substituted gpV while decreased triggering is observed for the 61DDED64/AAAA variant. Expression of gpV and its derivatives was induced by 0.2% arabinose. (**F-J**) Binding of gpVN^1-160^ protein (I) as well as of gpV^48-78^ peptide variants (F-H, J) to HigC as monitored by ITC. The stoichiometry is calculated per HigC protein chain. (**K**,**L**) ΔHDX between HigC vs HigC:gpV^48–78^ (K) as well as gpV^48–78^ vs HigC:gpV^48–78^ (L) plotted as a heat map. (**M**) HigC4:gpV^48–78^ AlphaFold model coloured as a function of the ΔHDX. The gpV^48–78^ fragment is outlined in bold. Only two subunits of the HigC tetramer were coloured for clarity.

To probe the possible physical interaction between gpV and HigBAC in the context of infection, we N-terminally FLAG-tagged HigC in the context of the full HigBAC system and performed immunoprecipitation followed by tandem mass spectrometry (IP-MS/MS) during infection with both wild-type λ*vir* and the λ*vir^escaspe_1^* mutant. We sampled at the 30 minutes post-infection time point, by which time late genes such as structural proteins are expected to be expressed^46^. gpV is dramatically enriched in HigC pulldown samples in case of wild-type λ*vir* as compared to samples generated using λ*vir^escaspe_1^* (**Figure 4C**). This suggests that the *gpV S54P* substitution allows λ*vir^escaspe^* to circumvent HigBAC defence by preventing the gpV recognition by HigC and consequent HigBAC activation.

Prediction of the complex of the gpVN region and the SecB^TA^ tetramer by AlphaFold-Multimer yielded a high-confidence model, providing structural insight into how HigBAC could be triggered through direct recognition of the tail protein by the HigC chaperone (**Figure 4D**, **Figure S6B**). The structure suggests that HigC prevents the folding of the N-terminal amino acid region of gpVN (aa 1-78), which, reminiscent of the ChAD:SecB^TA^ interaction, becomes linear and largely disordered. The residues constituting the β2-β3 loop as well as the flanking β-strands β2 and β3 are predicted to wrap around the chaperone, with aromatic residues W36 and W67 slotting into the same pockets of the chaperone that mediate the ChAD recognition. The two aromatic residues are spaced out by ≍30 residues and are located in a context similar to that of the ChAD SLiMs, i.e. preceded by an aspartic acid and followed by a polar residue, serine or threonine. Finally, the acidic patch D61DED64 is predicted to make extensive electrostatic interactions with the chaperone.

Co-expression of wild-type gpV – but not the S54P variant – induces HigBAC toxicity (**Figure 4E**). We used this reductionist system to probe our structural model through mutagenesis. Co-expression with gpV variants that are expected to compromise the interaction with HigC reveal differences in importance for activation. Activation is still seen with W36A and – to some extent – disruption of the acid patch D61DED64. S54P and W67A variants on the other hand do not activate HigBAC toxicity. Importantly, the N-terminal residues 1-78 of gpVN fused with a stabilising C-terminal SUMO tag activates the TAC toxicity more efficiently than the full-length gpV, with the S54P substitution in gpV^1-78^-SUMO merely decreasing its activity, but not abolishing it fully as observed in the case of full-length gpV.

### gpV co-translationally competes with HigA ChAD for binding to HigC

Our structural modelling suggests the topology of the fully folded gpV is incompatible with the N-terminal region forming a complex with HigC. This is due to the β strands β2 and β3 that flank the disordered loop β2-β3 coming together to form part of a twisted β sheet^45^ (compare **Figure 4B** to **Figure 4D**). Supporting this, the C-terminally truncated and SUMO- tagged gpVN fragment (1-78) lacking the β-sheet element is a more potent activator of HigBAC than the full-length gpV (**Figure 4E**). Since gpV in the cell readily folds and assembles into a tail tube superstructure, assisted by phage-encoded chaperones gpG and gpGT^47^, we reasoned that there is only a short window of opportunity for HigC to recognise the nascent N-terminal fraction of gpV as the protein is being synthesised and is not yet folded.

To probe the interaction between unfolded gpV and HigC as would occur in a co-translational context, we designed an unstructured gpV peptide fragment comprising residues 48-78 that contains the W67 and D61DED64 elements predicted to be recognised by HigC. ITC experiments showed that the two partners form a tight complex (K*D* = 250 nM, n = 0.47) (**Figure 4F**, **Table 1**). The W67A substitution disrupts the interaction between the gpV^48–78^ peptide, while the poly-alanine substitution of the D61DED64 element decreases the affinity more than 20-fold (K*D* = 7 μM, n = 0.5; n is calculated per HigC monomer) (**Figure 4G**,**H**). In contrast, our ITC experiments detect no interaction between gpVN (1-160) and HigC, even when the phage protein is used at 350 μM concentration and HigC at 30 μM, a 5-fold increase compared to the concentrations used for the peptide-HigC interactions (**Figure 4I**). The lack of interaction further supports the proposed requirement of gpV to be incompletely folded for gpV:HigC complex formation to take place. To mimic HigBAC triggering by gpV, we performed a competition experiment with the ChAD of HigA. HigC was saturated with 1.5- fold molar excess of the HigA^121–148^ ChAD-mimicking peptide, followed by titration with the gpV^48–78^ peptide while keeping the ChAD peptide concentration stable. Under these conditions gpV^48–78^ binds HigC with an effective K*D* of 6.6 μM, an about 25-fold drop in affinity (**Figure 4J**), which is strongly suggestive of direct competition between the two peptide ligands for HigC.

Finally, we probed the HigC:gpV interaction through hydrogen deuterium exchange mass spectrometry (HDX-MS) (**Figure 4K-M**, **Figure S5C**). To map the binding interface, we compared the deuterium exchange – a proxy for solvent accessibility – for individual HigC and gpV^48–78^ with that for the HigC:gpV^48–78^ complex (**Figure 4K-L**). A strong decrease in deuterium exchange (ΔHDX) in the HigC:gpV^48–78^ complex as compared to unbound HigC was localised to β1, β2, β3 and α1 elements of HigC that constitute the peptide binding groove as well as the pockets that accommodate the aromatic residues of the ChAD (**Figure 4M**). On the gpV^48–78^ side, the entire peptide was strongly protected from deuterium exchange (**Figure 4M**). These results are in excellent agreement with AlphaFold modelling and functional assays.

### A hybrid TAC system composed of the HigC chaperone and the CmdTA TA unit has an expanded antiphage defence spectrum and increased potency

Earlier studies have shown that SecB^TA^ chaperones are strictly specific and can only recognise the ChAD elements of cognate TA units^18^. To establish whether this is also the case for our two TAC systems, we tested the ability of non-cognate SecBs^TA^ – as well as that of housekeeping *E. coli* SecB – to neutralise the toxicity of HigBA and CmdTA when expressed *in trans* (**Figure 5A**). As expected, the non-cognate CmdC SecB^TA^ fails to counter the HigBA toxicity. Surprisingly however, the HigC chaperone neutralises the CmdTA unit efficiently, displaying better neutralising activity than the cognate chaperone CmdC. Overexpression of the housekeeping *E. coli* SecB chaperone partially suppresses the toxicity of both HigBA and CmdTA. In the case of HigBA, the housekeeping *E. coli* SecB is more efficient than the non- cognate CmdC SecB^TA^ chaperone. We have observed analogous results when hybrid TAC systems were cloned as one operon (**Figure S2**), suggesting the failure of CmdC to neutralise HigBA is not due to the disruption of co-translational folding of the TAC complex but reflects the intrinsic differences in the two SecB^TA^s.

**Figure 5.**
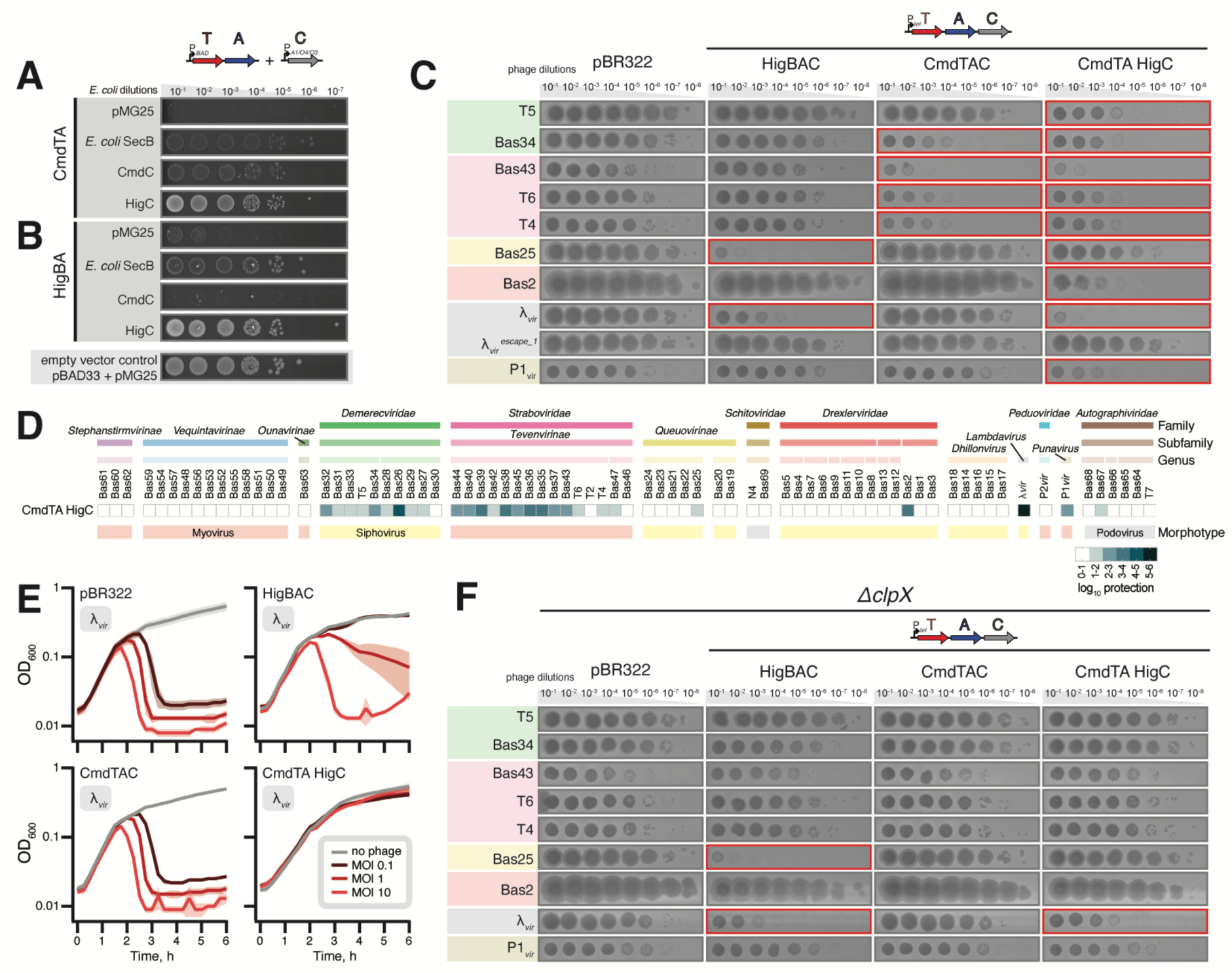
Engineering of a synthetic antiphage defence system through combination of the CmdTA toxin-antitoxin unit and the HigC phage-sensing chaperone. (**A**,**B**) Suppression of CmdTA (A) or HigBA (B) toxicity through IPTG-induced co-expression of SecB chaperones: *E. coli* housekeeping SecB, CmdC and HigC; pMG25 was used as an empty vector control. Expression TA and chaperone was induced by 0.2% arabinose and 500 µM IPTG, respectively. (**C**,**D**) *E. coli* BW25113 strains expressing HigBAC, CmdTAC or the hybrid CmdTA-HigC TAC operon composed of CmdTA toxin-antitoxin unit and HigC chaperone were challenged with ten-fold serial dilutions of BASEL^22^ and common lab coliphages, including the λ*vir^escape_1^* escape mutant variant. Panel (C) depicts results with select phages and the results of the full screen are shown in panel (D). (**E**) Growth of *E. coli* BW25113 carrying the empty vector or the indicated plasmid-encoded TAC systems in the presence of λ*vir* at MOIs of 0, 0.1, 1 and 10. (**F**) Activity of HigBAC, CmdTAC and the hybrid TAC in antiphage immunity tested in Δ*clpXE. coli*.

The ability of HigC to recognise and neutralise the non-cognate CmdTA makes it possible to directly test the functions of the TA unit and the chaperone in sensing the phage infection. We cloned the hybrid TAC system comprised of CmdTA followed by the HigC chaperone for constitutive expression from a pBR322 derivative plasmid and performed an immunity screen with the BASEL phage collection. Our initial naïve expectation was that the nature of the SecBs^TA^ chaperone (HigC in this case) would determine the spectrum of conferred defence, as the spectrum of the recognised phage triggers is defined by the nature of the chaperone. However, the result was strikingly different (**Figure 5C**,**D**). The hybrid TAC has a defence spectrum that i) almost fully combines that of the parental systems, affording stronger protection than either of the parental systems for some phages (such as λ*vir* and phages in the *Tevenvirinae* subfamily), ii) expands the spectrum of defence to all the three phage morphotypes, defending against viruses that were not protected against by neither of the parental systems, such as additional myo- and siphoviruses (*Punavirus* P1*vir*, Bas2 in the *Drexleviridae* family and multiple phages in the *Demerecviridae* family, including T5) as well as podoviruses from the *Autographviridae* family (Bas66 and Bas67, weak protection). Importantly, the λ*vir^escape_1^* mutant variant efficiently overcomes the immunity mediated by the CmdTA_HigC system, suggesting that HigC-mediated phage sensing operates similarly in hybrid TAC as it does in the native HigBAC (**Figure 5C**). Liquid culture λ*vir* infection assays with increasing MOI (0.1, 1 and 10) are in good agreement with the plaquing experiments (**Figure 5E**). While CmdTAC fails to provide any protection against the phage, the hybrid CmdTA-HigC system renders *E. coli* growth insensitive to λ*vir*, even at the MOI of 10. The effect is reminiscent of the full protection granted by CmdTAC against Bas43 in liquid culture experiments (**Figure 1G**).

Finally, we tested the ability of HigBAC, CmdTAC and the hybrid TAC to confer antiphage immunity in Δ*clpX E. coli* (**Figure 5F**). In good agreement with toxicity assays (**Figure 3H**,**I**), CmdTAC loses its defensive activity in Δ*clpX* background while HigBAC is as active as in the wild-type *E. coli*. The hybrid CmdTA-HigC system is functionally compromised in Δ*clpX E. coli*, which is to be expected as it shares the ChAD element with the ClpPX-dependent CmdTAC.

## Discussion

The CmdT TAC toxin from *E. coli* O112ab:H26 provides an example of an ART enzyme targeting mRNA to inhibit protein synthesis. While ART effectors of type VI secretion systems are known to target RNA, these systems specifically modify structured RNA: *Photorhabdus laumondii* Rhs ADP-ribosylates the 23S rRNA Sarcin-Ricin Loop^48^ and *Pseudomonas aeruginosa* RhsP2 targets diverse structured non-coding RNAs such as tRNA, tmRNA, 4.5S rRNA and RNase P^49^. The first CmdT system to be discovered, PD-T4-9 from *E. coli* ECOR22^20^, was also shown to target mRNA (see the accompanying paper by Vassalo and colleagues), thus suggesting the generality of this mechanism.

This study has shed light on how the chaperone-mediated recognition of viral structures has been adopted by TAC defence systems to sense phage infection. Phage structural proteins provide pathogen-associated molecular patterns (PAMPs) that are recognised by diverse defence systems such as the CapRel^SJ^^46^ toxSAS fused TA system, Avs STAND NTPases and AbiT abortive infection system^42,43,50–53^. Here we show that HigBAC from *E. coli* strain NT1F31 senses λ phage through recognition of the major tail protein gpV by the HigC chaperone. We propose the following model for HigBAC triggering (**Figure 6**). In the absence of infection, the inactive TAC complex is co-translationally assembled, with the aromatic residues of the HigA ChAD nucleating ChAD recognition by the HigC SecB^TA^ chaperone. Expression of gpV diverts the holdase activity of HigC, which leads to inactivation of the HigA antitoxin, unleashing the HigB RNase toxin that restricts the productive phage replication.

**Figure 6.**
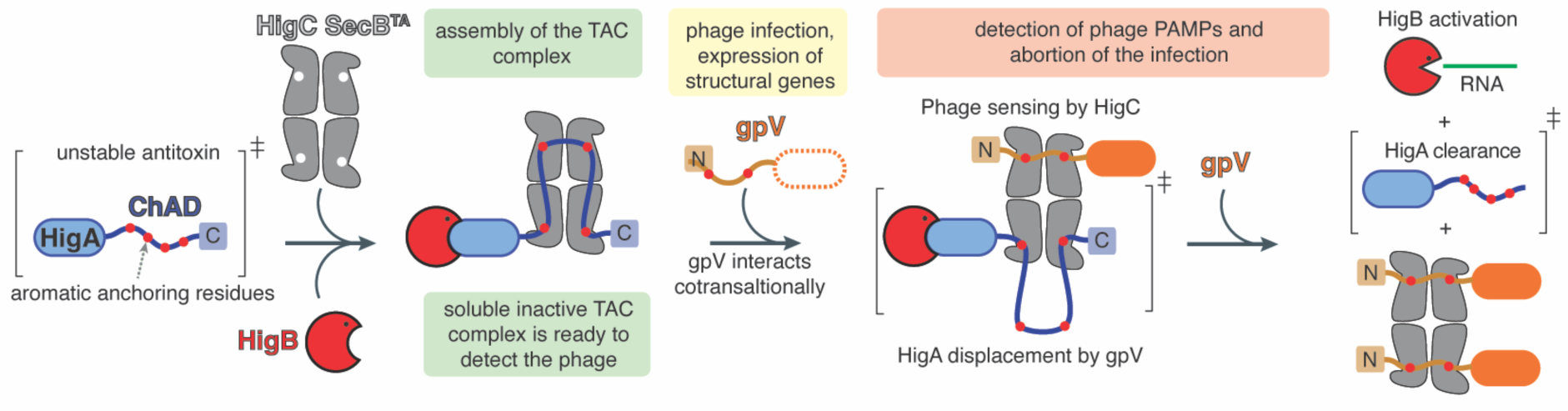
Sensing and abortion of phage infection by HigBAC defence system. In the absence of phage infection, the HigB RNase toxin, HigA antitoxin and tetrameric HigC4 chaperone form an inert complex. TAC complex formation stabilises otherwise unstable HigA. Both HigA destabilisation and HigA recognition are driven by the aromatic residues located in the ChAD region. Expression of the λ gpV protein is cotranslationally recognised by the HigC chaperone, resulting in direct competition between the HigA and the major tail protein. Destabilisation of HigA triggers HigB toxicity which, in turn, restricts the phage propagation in the infected cell.

Phage replication and assembly commonly relies on both host and phage-encoded chaperones. GroEL/GroES and DnaK/DnaJ/GrpE chaperone systems were discovered as essential components for λ replication in *E. coli* through genetic screens selecting for the *gro* (inability to *gro*w the phage) phenotype^54^. GroEL is essential for folding and assembly of phage structural elements^55^. The dedicated phage-encoded chaperones gpG and gpGT are essential for the assembly of the λ phage tail from gpV subunits^47,56^. In the phage T4, a dedicated co-chaperonin gp31 substitutes for GroES to facilitate the folding of the major capsid protein gp23^57,58^. Thus chaperones and their effects on modulating protein folding and aggregation are important facets of phage infection cycles, host:virus interactions, and – in the case of prophage-encoded defence systems such as TACs – virus:virus interactions.

### Limitations

This study lays the foundations for detailed studies of TAC-mediated antiphage defence. To uncover the molecular details, dedicated experimental structural studies are essential. Without these, understanding the molecular basis of SecB^TA^ specificity towards phage triggers and TAC antitoxin ChAD elements will remain inherently limited. The current study relies heavily on reconstitution of TAC sensing through co-expression of the identified phage trigger with the TAC system. Characterisation of the HigBAC TAC expression and assembly dynamics – and its subversion by the gpV – during the phage infection is essential for uncovering the co-translational nature of the SecB^TA^-based sensory system. Finally, dedicated substrate specificity studies are needed to uncover the specific RNA species and RNA motifs targeted by HigB and CmdC TAC toxins.

## ACKNOWLEDGMENTS

The AlphaFold2 computations were enabled by the supercomputing resources Berzelius provided by the National Supercomputer Centre (NSC) at Linköping University and the Knut and Alice Wallenberg foundation. Additional computational resources used were provided by the National Academic Infrastructure for Supercomputing in Sweden (NAISS) and the Swedish National Infrastructure for Computing at NSC, Chalmers University Centre for Computational Science and Engineering (C3SE), and PDC Centre for High Performance Computing, KTH Royal Institute of Technology, partially funded by the Swedish Research Council through grants 2018-05973 and 2022-06725. This work was supported by the Knut and Alice Wallenberg Foundation (project grant 2020-0037 to GCA and VH), the Swedish Research Council (Vetenskapsrådet) grants (2019-01085, 2022-01603 and 2023-02353 to GCA, 2021- 01146 to VH), Crafoord foundation (project grant Nr 20220562 to VH), the Estonian Research Council (PRG335 to VH and TT), Cancerfonden (20 0872 Pj to VH), the Fonds National de Recherche Scientifique (FNRS CDR J.0068.19 and J.0065.23F; FNRS-EQP UN.025.19; FNRS PDR T.0066.18 and PDR T.0090.22 to AG-P), ERC (CoG DiStRes, n° 864311 to AG- P), Fonds Jean Brachet and the Fondation Van Buuren (AG-P), FNRS-ASP (SZC) as well as Ambizione Fellowship PZ00P3_180085 and Starting Grant TMSGI3_211369 of the Swiss National Science Foundation (SNSF) (both AH). MTL is an Investigator of the Howard Hughes Medical Institute. JAB and HS were funded by BBSRC grants BB/X003035/1 and BB/T017570/1. K.C.W. is a FRIA fellow, C.M. is supported as a Research Associate of the FRS–FNRS. C.M. was supported by grant F.4532.22 from the FRS–FNRS.

## AUTHOR CONTRIBUTIONS

VH coordinated the study and drafted the manuscript with contributions from all authors. TM, TK, KE, MJOJ, SZC, GME, CRD, MTL, TT, AH, CM, HS, AG-P, GCA and VH designed experiments and analysed the data. TM, TK, KE, MJOJ, SZC, GME, LS, KCW, CM, JAB, CRD and TB performed experiments. RO, JAN, AAE, AG-P and GCA performed bioinformatic analyses.

## STAR ★ METHODS

Detailed methods are provided in the online version of this paper and include the following:

- KEY RESOURCES TABLE
- CONTACT FOR REAGENT AND RESOURCE SHARING
- EXPERIMENTAL MODEL AND SUBJECT DETAILS

`o` Bacterial strains
- METHOD DETAILS

`o` Figure preparation
`o` Bioinformatics
`o` Construction of plasmids
`o` Microbiological assays
`o` *In vivo* functional assays
`o` Biochemical assays
- QUANTIFICATION AND STATISTICAL ANALYSIS

`o` Statistical analysis of IP-MS/MS data
- DATA AND SOFTWARE AVAILABILITY

`o` The study does not make use of unpublished data or software

## STAR ★ METHODS

### Resource Availability

**Lead Contact and Material availability**

Please direct any requests for further information or reagents to the Lead Contact: Vasili Hauryliuk.

## EXPERIMENTAL MODEL AND SUBJECT DETAILS METHOD DETAILS

### Bacterial strains

Bacterial strains, bacteriophages, plasmids as well as oligonucleotide primers used in the study are listed in **Dataset S1**.

### Figure preparation

Figures were prepared using UCSF ChimeraX 1.6.1, GraphPad Prism 10.1.1 (GraphPad Software), Igor Pro 7.07 (WaveMetrics, Inc.), Adobe Illustrator 28.0 (Adobe Inc.) and Adobe Photoshop 28.0 (Adobe Inc.), CamSol^38^ and Fiji^59^.

### Bioinformatics

#### Genomic neighbourhood search and multiple sequence alignment

Sequence searching for SecB homologues was carried with PSI-BLAST^60^ against the RefSeq database, limiting by taxonomy to Enterobacteriaceae and an E value threshold of 0.01. The query was *Lactiplantibacillus garii* accession WP_125072952, identified with NetFlax as a SecB-like protein associated with a MqsRA-like TA^21^. Eight PSI-Blast iterations were carried out, with all identified hits going through to each round, after which no new hits were identified. All 558 hits were used as queries for FlaGs2^21^ (https://github.com/GCA-VH-lab/FlaGs2) with default settings to produce (**Dataset S1**). Chaperone and Antitoxin sequences were aligned using MAFFT L-INS-i^61^. Alignments were visualised with AliView^62^ and Jalview^63^. Prophage boundaries were predicted with the PHASTER^64^ server.

#### Structural modelling

Protein structure predictions were made with AlphaFold2 v2.3.1^35^ and complexes were predicted with the AlphaFold-Multimer^34^ protocol. The structural template cut-off date was set to May 14, 2020 (--max_template_date=2020-05-14). The quality of structural predictions is assessed with predicted template-modelling (TM) scores^65^; pTM scores all residues and ipTM scores interfacing residues. Models with the best pTM+ipTM scores were chosen for analysis. pTM and ipTM scores of 0.5 are indicative of correct folds, while scores of 0.8 are indicative of high quality models^33^. HigA:HigB:HigC4 and gpVN:HigC4 models have pTM+ipTM scores of 0.724 and 0.740 respectively.

#### Phage clustering based on proteome similarity

BASEL phages were clustered through hierarchical clustering using pairwise proteome similarity scores. All proteins from the BASEL collection of phages were clustered using MMseqs^66^ with optional parameters (*--cluster-mode 1 --cov-mode 0 -c 0.7 --min-seq-id 0.3*), with each protein being assigned to a cluster corresponding to a set of homologues. To calculate the similarity scores between the i^th^ and j^th^ phages (*sim(i,j)*) the number of overlapped homologues was normalized to the size of the i^th^ proteome. A symmetric proteome composition distance (PCD, *dist(i,j)*) matrix was calculated with the formula *dist(i,j) = 1 – (average(sim(i,j), sim(j,i)))*. The clustering was performed in R using the average-linkage method and data were visualised using the *ComplexHeatmap* library^67^. Phages were annotated with family, subfamily and genus using taxonomic information extracted from the NCBI Taxonomy database^68^ (January 2024 version) and ICTV^69^ (2022 update).

### Construction of plasmids

SnapGene (GSL Biotech LLC) and Geneious Prime (Biomatters) software were employed for primer and plasmid map design. All plasmids were constructed with either circular polymerase extension cloning (CPEC)^70,71^ with Phusion polymerase (Thermo Scientific), Gibson assembly^72^, or ligation of DNA fragments with T4 DNA Ligase (Thermo Scientific). The following TAC genes were used: CmdT WP_097333619.1, CmdA WP_053886482.1, CmdC WP_053886483.1, HigB WP_112844290.1, HigA WP_112844289.1 and HigC WP_112844288.1. pBAD33 plasmid (arabinose inducible PBAD promoter) was used to express TA and TAC regions and their variants, pAS004, a pBAD33 derivative, was used to express λ*vir* gpV and its variants, and pMG25 (IPTG inducible PA1/O4/O3 promoter) vector was used for chaperone expression. The HigBA, HigBAC, and their mutated version fragments were inserted into the pBAD33 plasmid with a weak Shine-Dalgarno motif (ATTAGAAGAATAAG) from CmdTAC. pAS004, pMG25, and CmdTAC pBAD33 derivates were created with a strong SD (AGGAGGAATTAA). For immunity assays, TAC regions (coding sequences with 150 bp upstream for HigBAC region, and 118 bp upstream for CmdTAC region and native terminators for both predicted by ARNold^73^) were cloned into the pBR322 vector under the control of the constitutive Ptet promoter. For metabolic labelling assays the TAC toxin genes were cloned into pBAD33, either without the Shine-Dalgarno sequence (the more toxic HigB, VHp1718) or with a strong Shine-Dalgarno sequence AGGAGGAATTAA (the less toxic CmdT, VHp1140). pBR322-HigBA/FLAG-C construct used for pulldown experiments was created by site-directed mutagenesis using pBR322- HigBAC as a template. Detailed cloning schemes can be found in **Dataset S1**. All constructs were verified by sequencing.

### Microbiological assays

#### TA and TAC toxicity assays

The TA regions of *E. coli* NT1F31 HigBAC system and *E. coli* O112ab:H26 CmdTAC system were expressed from pBAD33. HigBA and its derivatives were expressed with a weak Shine-Dalgarno motif from CmdTAC (ATTAGAAGAATAAG), while CmdTA with a strong Shine-Dalgarno sequence (AGGAGGAATTAA). Chaperones HigC, CmdC, and *E. coli* BW25113 housekeeping chaperone SecB were expressed with a strong SD from pMG25 plasmid. Plasmids were transformed into wild-type *E. coli* BW25113 or protease deletion strains. Bacterial cultures from single colonies were grown for five to six hours, adjusted to OD600 1.0, serially diluted (10^1^- to 10^8^-fold) and spotted on LB medium supplemented with 100 μg/mL carbenicillin (Fisher Bioreagents), 25 μg/mL chloramphenicol (AppliChem) and inducers (0.2% arabinose for TA induction, 50 µM or 500 µM IPTG for chaperone induction). Plates were scored after overnight incubation at 37 °C. For *in cis* TAC neutralization assays the CDSs of HigBAC or CmdTAC were expressed from pBAD33, with a weak SD for HigBAC and a strong SD for CmdTAC. Plasmids expressing the TA or TAC region were transformed into *E. coli* BW25113. Bacterial cultures were started from single colonies, grown for five to six hours, adjusted to OD600 1.0, serially diluted (from 10^1^- to 10^8^-fold) and spotted on LB medium supplemented with 25 μg/mL chloramphenicol (AppliChem) and 0.2% arabinose for induction. Plates were incubated overnight at 37 °C and scored.

#### TAC triggering assays

The effects of co-expression of *E. coli* NT1F31 HigBAC (pBR322 with constitutive Ptet-driven expression) and λ*vir* gpV variants (pAS004 with arabinose-inducible PBAD-driven expression) were tested on a spot assay. The plasmids were transformed into *E. coli* BW25113 and bacterial cultures from single colonies were grown for five to six hours. Optical density was adjusted to OD600 1.0, serially diluted, and spotted on LB medium supplemented with 100 μg/mL carbenicillin (Fisher Bioreagents), 20 μg/mL gentamicin (Sigma-Aldrich) and 0.2% arabinose for gpV induction. Plates were scored after an overnight incubation at 37 °C.

#### Experimental phage infections

To assess the effect of the TAC systems on phage defence, we performed efficiency of plating assays, essentially as described previously^21^, using the BASEL collection^22^ and a subset of common laboratory phages (**Dataset S1**). Briefly, overnight cultures of *E. coli* BW25113 cells carrying either an empty vector (pBR322 derivative lacking the tetracycline resistance cassette^30^) or a TAC system (pBR322-Ptet-HigBAC (VHp1259), pBR322-Ptet-CmdTAC (VHp1256), or pBR322-Ptet-CmdTA-HigC (VHp1608)) were mixed with top agar (LB with 0.5% agar, 20 mM MgSO4, and 5mM CaCl2), to a final concentration of 0.075 OD600 units/ml, and overlayed on LB-agar plates (1.5% agar). Individual phage stocks were 10-fold serially diluted in SM buffer (0.1 M NaCl, 10 mM MgSO4, and 0.05 M Tris-HCl pH 7.5) and 2.5 μL of each of eight dilutions spotted on solidified top agar plates. Plaque formation was monitored after 6 and 24h of incubation at 37 °C. The phages that showed sensitivity to at least one system were re-tested towards all three systems using three different transformants for each plasmid. Plaques were counted after 24 h and the efficiency of plaquing (EOP) was determined for each repeat by dividing the plaque forming units for a given TAC system by that for the vector control. The average EOP value from the three repeats was -log10 transformed, yielding the log10 protection value used in heatmaps.

For phage infection of liquid cultures, three different transformants of *E. coli* BW25113 harbouring either an empty vector or a TAC-containing plasmid were grown overnight in LB medium supplemented with ampicillin, 10 mM MgSO4, and 2.5 mM CaCl2. The cells were diluted to OD600 ≍ 0.075 in the same medium and 100 μL were added to wells of a 96-well plate. The relevant phages were diluted in SM buffer to generate final MOI values of 10, 1, and 0.1 when 10 μL of the dilution was added to a well. Ten μL of SM buffer were added to control wells. The growth was monitored at 37 °C in a Synergy H1 (BioTek) plate reader measuring OD600 every 15 min.

#### Isolation of bacteriophage escape mutants

To identify bacteriophage escape mutants, 200 µl of overnight culture of *E. coli* BW25113 strain carrying pBR322-Ptet-HigBAC (VHp1259) plasmid was infected with 20 µl of different λ*vir* stock dilutions, mixed with top agar and poured on LB agar plate. After incubation at 37 °C for 24h, plaques with normal morphology (as seen on control culture without the immunity system) were isolated with a sterile toothpick and re-streaked three times using BW25113 carrying pBR322-Ptet-HigBAC plasmid as the host. High-titer stocks were prepared from potential escape mutants as described previously^22^. Norgen Biotek Phage DNA Isolation Kit (Norgen Biotek Corp.) was used to isolate genomic DNA of bacteriophages. The DNA was sequenced at the Microbial Genome Sequencing Center and results analysed using Geneious Prime (Biomatters).

### *In vivo* functional assays

#### Metabolic labelling with ^35^S-methionine, ^3^H-uridine and ^3^H-thymidine

For metabolic labelling experiments the *E. coli* BW25113 strain was co-transformed with the pBAD33 plasmid carrying the toxin gene of interest (CmdT:VHp1140 and HigB:VHp1718) for L-arabinose-inducible expression and the empty pMG25 vector. Transformed cells were initially plated on LB plates supplemented with 100 μg/mL carbenicillin, 25 μg/mL chloramphenicol, and 0.2% glucose (to suppress leaky toxin expression). Using individual *E. coli* colonies for inoculation, 2 mL liquid cultures were prepared in defined Neidhardt MOPS minimal media^74^, supplemented with 100 μg/mL carbenicillin, 25 μg/mL chloramphenicol, 0.1% casamino acids, and 0.2% glucose, and grown overnight at 37 °C with shaking. Subsequently, experimental 15 mL cultures were prepared in 125 mL conical flasks in MOPS medium, supplemented with 0.5% glycerol, 100 μg/mL carbenicillin, 25 μg/mL chloramphenicol, as well as a set of 19 amino acids (lacking methionine), each at a final concentration of 25 μg/mL. These cultures were inoculated overnight to reach a final OD600 of 0.05 and grown at 37 °C with shaking until the OD600 reached 0.2. At this point, 1 mL aliquots (designated as the pre-induction zero time-point) were transferred to 1.5 mL Eppendorf tubes containing 10 μL of the respective radioisotope (^35^S methionine - 4.35 µCi, Hartman; ^3^H uridine - 0.65 µCi, Hartman; or ^3^H thymidine - 2 µCi, Hartman) and placed in a heat block at 37 °C. Toxin expression in the remaining 14 mL culture was induced by adding L-arabinose to a final concentration of 0.2%. Throughout the toxin induction time course, 1 mL aliquots were taken from the 15 mL culture and transferred to 1.5 mL Eppendorf tubes containing 10 μL of the appropriate radioisotope (^35^S methionine, ^3^H uridine, or ^3^H thymidine). Radioisotope incorporation was halted after 8 minutes of incubation at 37 °C by adding 200 μL of ice-cold 50% trichloroacetic acid (TCA) to the 1 mL cultures. Additionally, 1 mL aliquots were periodically sampled for OD600 measurements. The resultant 1.2 mL culture/TCA samples were loaded onto GF/C filters (Whatman) prewashed with 5% TCA and unincorporated label was removed by washing the filter twice with 5 mL of ice-cold TCA followed by a 5 mL wash with 95% EtOH (twice). The filters were placed in scintillation vials, dried for at least two hours at room temperature, followed by the addition of EcoLite™-scintillation cocktail (5 mL per vial; MP Biomedicals). After shaking for 15 minutes, radioactivity was quantified using Tri-Carb 4910TR-scintillation counter (Perkin Elmer). Isotope incorporation was quantified by normalizing radioactivity counts (CPM) to OD600, with the pre-induction zero time-point serving as the reference (set to 100%). All experiments were conducted in triplicates, using three independent cultures initiated from distinct colonies.

#### Immunoblotting

*E. coli* BW25113 strain co-transformed with pBAD33 plasmid carrying either the N-terminally His6 tagged *higA* gene (VHp1720) or its tetra-substituted version [4*: *W128A F158A F188A Y214A*] (VHp1722) for L-arabinose-inducible expression, together with either empty pMG25 vector (VHp1069) or its derivative for IPTG-inducible expression of HigC (VHp1618). Experimental 20 mL cultures (LB media supplemented with 100 μg/mL carbenicillin, 25 μg/mL chloramphenicol and 50 µM IPTG) were inoculated to an OD600 of 0.05 and grown at 37 °C with shaking until reaching the OD600 of 0.5. Expression of His6HigA was induced by addition of L-arabinose to a final concentration of 0.2%, 1 mL samples were collected after 1 hour, the cells pelleted by centrifugation, dissolved in 1x SDS-PAGE sample buffer (200 μL for 1 mL of OD600 1.0 culture), denatured at 95 °C for 5 min and 10 μL of the final sample were resolved on SDS-PAGE (12% acrylamide/bis-acrylamide 37.5:1). Proteins were transferred to BioTrace™ NT nitrocellulose membranes (Pall Life Sciences) using the Trans- Blot^®^ Turbo^TM^ Transfer System (Bio-Rad), and the membranes were blocked for 1 hour in PBS-T with 5% skimmed milk at room temperature. Blocked membranes were incubated with either primary anti-His-tag antibodies (Boster Biological Technology, M30975; 1∶1000) or primary anti-RpoB antibodies (Abcam, ab191598; 1:2000) at 4 °C overnight in PBS-T with 1% milk. After three 5-minute washes with fresh PBS-T, HRP-conjugated secondary antibodies were added (goat anti-mouse IgG, Argisera, AS11 1772; 1:5000 or anti-rabbit IgG, Sigma-Aldrich, A0545; 1:5000, respectively; both diluted in PBS-T) and the membranes were incubated for 1 hour at room temperature. The membranes were washed twice with PBS-T for 5 minutes followed by one 5-minute wash with PBS. The WesternBright Quantum HRP substrate (Advansta) signal was visualized with Amersham™ ImageQuant 800 (Cytiva) imaging system.

#### Fluorescence microscopy

Fluorescence microscopy was carried out with early-mid logarithmic growth phase *E. coli* BW25113 cells grown in LB media (Miller) (10 g l^−1^ tryptone, 5 g l^−1^yeast extract, 10 g l^−1^ NaCl) supplemented with 100 µg/ml ampicillin for maintaining plasmids at 37 °C. Samples were immobilized on Teflon-coated multi-spot microscope slides (Thermo Fisher) covered with a thin layer of H2O/1.2 % agarose and imaged immediately. Microscopy was performed using a Nikon TI2 equipped with Nikon CFI Plan Apo DM Lambda 100X Oil objective, CoolLED pE-4000 light source, and Photometrics Kinetix sCMOS camera. Images were acquired with Nikon NIS-Elements AR software and analysed with Fiji.^59^ *Immunoprecipitation followed by tandem mass spectrometry (IP-MS/MS)*

Overnight cultures containing plasmid-based anhydrous tetracycline (aTc)-inducible HigBAC or HigBA/FLAG-C cells were back-diluted in 250 mL LB with 0.05 mg/mL carbenicillin and 100 ng/mL aTc and grown at 37 °C to an OD600 = 0.2. Cultures were infected with either λ*vir* or λ*vir^escape_1^* mutant at an MOI 10 and samples collected at 0-, 15-, and 30-minutes post-infection. To collect samples, the culture was pelleted at 7,500g for 5 minutes, the pellet decanted and then resuspended in lysis buffer (25 mM Tris-HCL, 150 mM NaCl, 1 mM EDTA, 5% glycerol, 1% Triton X100) supplemented with 1 μL/mL Ready-Lyse™ Lysozyme (Fischer Scientific), 1 μL/mL benzonase (Sigma), and cOmplete™ Protease Inhibitor Cocktail (Roche) and then flash frozen in liquid nitrogen. Samples were thawed and additional lysis buffer added as necessary to normalize sample concentration by OD600 value taken concurrent with sample collection. Samples were refrozen in liquid nitrogen and thawed to ensure complete cell lysis. Samples were spun at 20,000g for 10 minutes at 4 °C to pellet any debris. For each sample, 50 μL of Pierce™ Anti-DYKDDDDK magnetic agarose beads was mixed with 450 μL of lysis buffer and then collected to the side of the tube using a magnetic rack. Beads were then washed twice with 500 μL of lysis buffer. After the final wash, beads were mixed with 1 mL of sample and incubated for 20 minutes at room temperature on an end-to-end rotor. After incubation, beads were washed with wash buffer (1X PBS, 150 mM NaCl) twice and then once with MiliQ H2O. On-bead reduction, trypsin digest, and LC-MS/MS were done as previously^42^. Detected peptides were mapped to MG1655 and λ protein sequences and the abundance of proteins were estimated by number of spectrum counts/molecular mass to normalize for protein sizes.

### Biochemical assays

#### In vitro translation assays

PURExpress *in vitro* protein synthesis kit (NEB, E6800) supplemented with 0.8 U/µL RNase Inhibitor Murine (NEB, M0314S) was used for reactions as per the manufacturer instructions. All reactions contained each ART template plasmids (10 ng/µL, VHp1221) with or without 0.1 mM NAD^+^ or 6-biotin-17-NAD^+^. After a 10-minute incubation at 37 °C, a 1.3 µL aliquot of the reaction mixture was taken and quenched by addition of 13.7 µL of 2x SDS-PAGE sample buffer (100 mM Tris:HCl pH = 6.8, 4% SDS, 0.02% bromophenol blue, 20% glycerol, 20 mM DTT and 4% β-mercaptoethanol), and DHFR template plasmid was added to the remaining reaction mixture at a final concentration of 20 ng/µL. After further incubation at 37 °C for 1 hour, the reaction mixture was mixed with 9-fold volume of 2x sample buffer, denatured at 95 °C for 5 min and resolved on SDS-PAGE gel (18% acrylamide/bis-acrylamide = 37.5:1). The SDS-PAGE gel was fixed by incubating for 5 min at room temperature in 50% ethanol solution supplemented with 2% phosphoric acid, washed three times with water for 20 min at room temperature, and stained with “blue silver” solution (0.12% Brilliant Blue G250 (Sigma-Aldrich, 27815), 10% ammonium sulfate, 10% phosphoric acid, and 20% methanol) overnight at room temperature. After washing with water for 3 hours at room temperature, the gel was imaged on an Amersham™ ImageQuant 800 (Cytiva) imaging system.

#### Immunoblotting

For streptavidin blotting, the PURExpress reaction was incubated with 0.1 mM 6-biotin-17- NAD^+^ and 5 nM PCR fragment amplified with VTK53 and VTK54 primers and VHp1221 template at 37 °C for 2 hours. In the case of model-substrate modification, 0.1 volume of 100 µM oligo RNA or DNA was added to the reaction at 30 min after starting the reaction then the reaction was incubated at 37 °C for a further 30 min. The sample was denatured in 1x SDS-PAGE sample buffer (for protein) at 95 °C for 5 min or in Urea-PAGE sample buffer (98% formamide, 10 mM EDTA, 0.3% BPB and 0.3% Xylene cyanol, for nucleic acids) at 65 °C for 5 min, and resolved on SDS-PAGE (12% acrylamide/bis-acrylamide = 37.5:1) or 8 M Urea-PAGE (6% or 8% acrylamide/bis-acrylamide = 19:1). When using double-strand model substrates, the sample was mixed with 0.2 volume of TriTrack DNA Loading Dye (6X)

(Thermo Scientific, R1161) and resolved on native PAGE (8% acrylamide/bis-acrylamide = 19:1). Resolved samples were transferred to Zeta-Probe^®^ Blotting Membranes (Bio-Rad) using Trans-Blot^®^ Turbo^TM^ Transfer System (Bio-Rad). The membrane was blocked in PBS with 3% BSA at room temperature for 30 min, and the first antibody incubation was performed at 4 °C for one hour in PBS with 3% BSA and HRP-conjugated streptavidin (Thermo scientific, N100; 1:5,000 dilution). After two 5-minute washes with fresh PBS-T and one 5-minute wash with PBS, biotinylated ADP signal was imaged on an Amersham™ ImageQuant 800 (Cytiva) imaging system using WesternBright Quantum HRP substrate (Advansta).

#### Purification of HigC

N-terminally His6-TEV-tagged HigC was cloned in pET28b vector and expressed in *E. coli* BL21(DE3). Cultures were grown in LB medium supplemented with kanamycin (50 μg/ml) at 37 °C with aeration. Expression was induced with 0.1 mM IPTG when the cells carrying the plasmid reached an OD600 nm of ∼ 0.5-0.8 at 25 °C. After induction, the cells were harvested 16h later by centrifugation and resuspended with buffer (25 mM HEPES, pH 7.6, 300 mM NaCl, 300 mM KCL and 2 mM MgCl2, 1mM TCEP) supplied with complete protease-inhibitor cocktail (Roche). The resuspended cells were flash-frozen in liquid nitrogen and stored at – 80 °C. The cell extracts were lysed using an Emulsiflex cell disruptor and the lysate was centrifuged to remove cell debris for 45 min at 25 000g. Prior to purification the extract was filter through a 0.45 μm membrane and loaded onto 1 ml HiTrap Ni NTA column (Cytiva) coupled to an FPLC (ÄKTA Explorer) equilibrated with buffer A (25 mM HEPES, pH 7.6, 300 mM NaCl, 300 mM KCL and 2 mM MgCl2, 1mM TCEP). The column was washed with buffer A before proceeding with the elution of HigC with buffer B (25 mM HEPES, pH 7.6, 300 mM NaCl, 300 mM KCl and 2 mM MgCl2, 1mM TCEP and 500 mM Imidazole). After tag removal by incubating the protein sample for 10h with TEV (1:5 molar ratio) at 10 °C the solution was passed though an HiTrap Ni NTA column to trap the protease and the His-tag and collect the tagless HigC. A final step of size exclusion chromatography was performed with the pooled samples in 25 mM HEPES, pH 7.6, 300 mM NaCl, 300 mM KCl and 1mM TCEP.

#### Purification of gpVN

N-terminally His6-TEV-tagged gpVN was cloned in pET28b vector and expressed in *E. coli* BL21(DE3). Cultures were grown in LB medium supplemented with kanamycin (50 μg/ml) at 37 °C with aeration. Expression was induced with 0.5 mM IPTG when the cells carrying the plasmid reached an OD600 nm of ∼ 0.5-0.8 at 25 °C. Cells were harvested by centrifugation 4h after induction and resuspended in 25 mM HEPES, pH 7.5, 300 mM NaCl, 300 mM KCl and 10 mM MgCl2, 1mM TCEP supplied with complete protease-inhibitor cocktail (Roche). DNase (10 μg/ml) was added to the resuspended cells before they were stored at –80 °C. The cell extracts were lysed using an Emulsiflex cell disruptor and the lysate was centrifuged to remove cell debris for 45 min at 42 000 g. The extract was filtered through a 0.45 μm membrane, then loaded onto a gravity-flow column (Cytiva) packed with Co^2+^-affinity resin equilibrated with purification buffer (25 mM HEPES, pH 7.5, 300 mM NaCl, 300 mM KCl 1mM TCEP). The column was washed with 10 ml purification buffer, then the protein was eluted stepwise with purification buffer containing 500 mM imidazole. The eluted fractions were transferred to a size-exclusion chromatography (SEC) column Superdex 75 pg (Cytiva), equilibrated in the SEC buffer (25 mM HEPES, pH 7.5, 150 mM NaCl, 150 mM KCl, 1mM TCEP). The His-TEV tag was removed by incubating with TEV enzyme (1:50 molar ratio) at 10 °C overnight. The cleaved protein was recovered by passing the sample over a gravity-flow column (Cytiva) packed with Co^2+^-affinity resin (Thermo Fisher Scientific).

#### Isothermal titration calorimetry (ITC)

All titrations were performed with an Affinity ITC (TA instruments) at 30 °C in 25 mM HEPES, 150 mM KCl, 150 mM NaCl, 1 mM TCEP at pH 7.5 (ITC buffer). The ChAD fragments and the gpV synthetic truncates were directly resuspended in the ITC buffer. The final concentrations were verified by the OD280 absorption using a Nanodrop One (ThermoScientific). All ITC measurements were performed by titrating a constant volume of 2 μl of peptide or gpVN into the ITC cell (containing HigC at 15 μM) using a constant stirring rate of 75 rpm. All data were processed, buffer-corrected and analyzed using the NanoAnalyse and Origin software packages. Sequences of synthetic polypeptides used in the study are provided in the (**Dataset S1**).

#### HDX-MS sample preparation

The HigC protein was concentrated to 100 μM. To prepare the complex of HigC with gpV^48–78^, the peptide was directly dissolved in the HigC sample in a 1:4 molar ratio. For each experiment, 10 μL of protein sample (HigC or HigC:gpV^48–78^ were incubated for 1 min, 5 min, <u>15 min or 60 min in 50 μL of labelling buffer (25 mM HEPES pH 7.5, 400 mM KCl, 400 mM</u> <u>NaCl, 1 mM TCEP) at 20</u> °C<u>. The non-deuterated reference points were prepared by replacing</u> <u>the labelling buffer with the equilibration buffer (25 mM HEPES pH 7.5, 400 mM KCl, 400</u> <u>mM NaCl, 1 mM TCEP). After labelling, the samples were quenched by mixing 60 μL of pre-chilled quench buffer (1.2 % formic acid, pH 2.4) with the labelled sample,</u> flash-frozen in liquid N2 and stored at -80 °C.

#### HDX-MS data collection

Prior to each injection, samples were thawed at room temperature and 150 µL of the quench samples were directly transferred to the Enzymate BEH Pepsin Column (Waters Corporation) at 200 µL per min and at 20 °C with a pressure of 3 kPSI. Peptic peptides were trapped for 3 min on an Acquity UPLC BEH C18 VanGuard Pre-column (Waters Corporation) at a 200 µL per min flow rate in water (0.1% formic acid in HPLC-grade water, pH 2.5) before elution to an Acquity UPLC BEH C18 Column for chromatographic separation. Separation was done with a linear gradient buffer (3–45% gradient of 0.1% formic acid in acetonitrile) at a flow rate of 40 µL per min. Peptides identification and deuteration uptake analysis was performed on the Synapt G2 in ESI ± MS^E^ mode (Waters Corporation). Leucine Enkephalin was applied for mass accuracy correction and sodium formate was used as calibration for the mass spectrometer. MS^E^ data were collected by a 20-30 V transfer collision energy ramp. The pepsin column was washed between injections using pepsin wash buffer (1.5 M Guanidinium HCl, 4% (v/v) acetonitrile, 0.8% (v/v) formic acid). A cleaning run was performed before loading the samples to prevent peptide carry-over. Optimized peptide identification and peptide coverage for all samples was performed from undeuterated controls (five replicates). All deuterium time points were performed in triplicates.

## QUANTIFICATION AND STATISTICAL ANALYSIS

Spectral counts for each protein in the *E. coli* MG1655 and λ*vir* proteomes and the HigBAC system were calculated, and high confidence hits assessed. The ratio of spectral counts between the λ*vir* and λ*vir^escape_1^* escape mutant infected samples at 30 minutes post-infection with a pseudocount added to each count was used to generate Figure 4C.

## DATA AND SOFTWARE AVAILABILITY

The study does not make use of unpublished data or software. The IP-MS mass spectrometry data to are deposited at the MassIVE database at doi:10.25345/C5T14V10M. AlphaFold2 predicted structures together with the accompanying quality metric scores are available in https://github.com/GCA-VH-lab/tac_ms_af. The phage annotation table, results of phage protein clustering including heatmaps, and the R script used for clustering are available at figshare [dx.doi.org/10.6084/m9.figshare.24968247].

## KEY RESOURCES TABLE

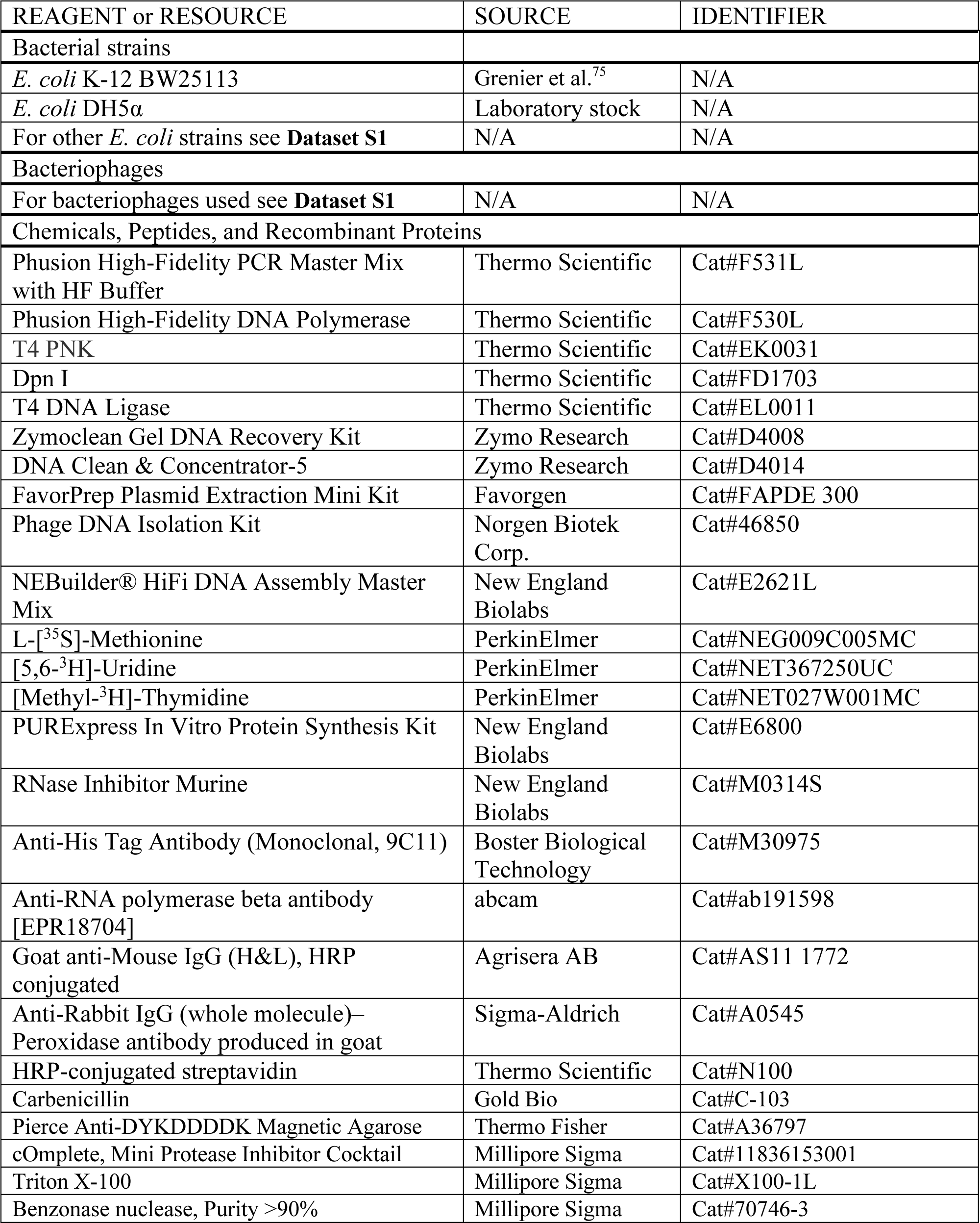

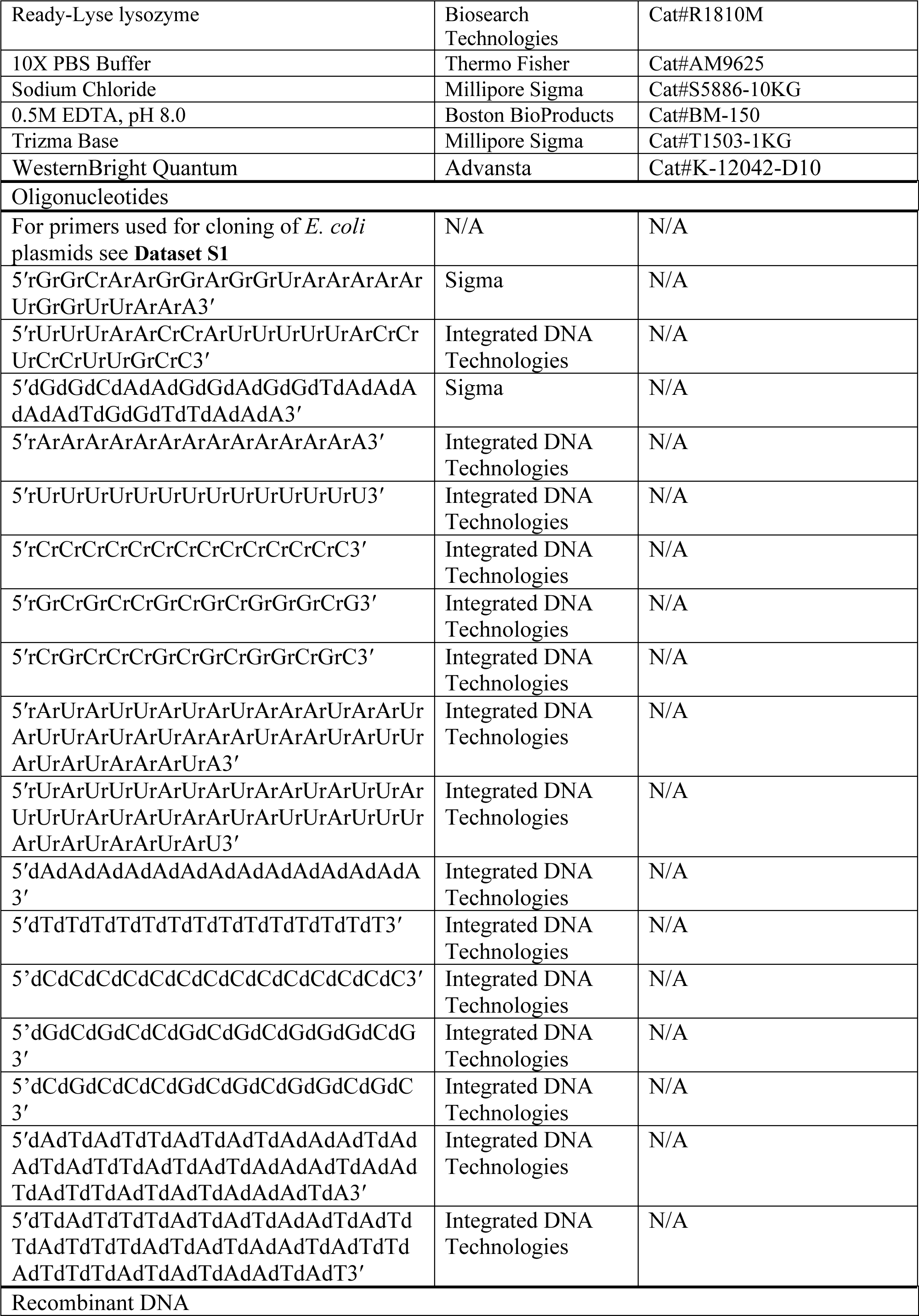

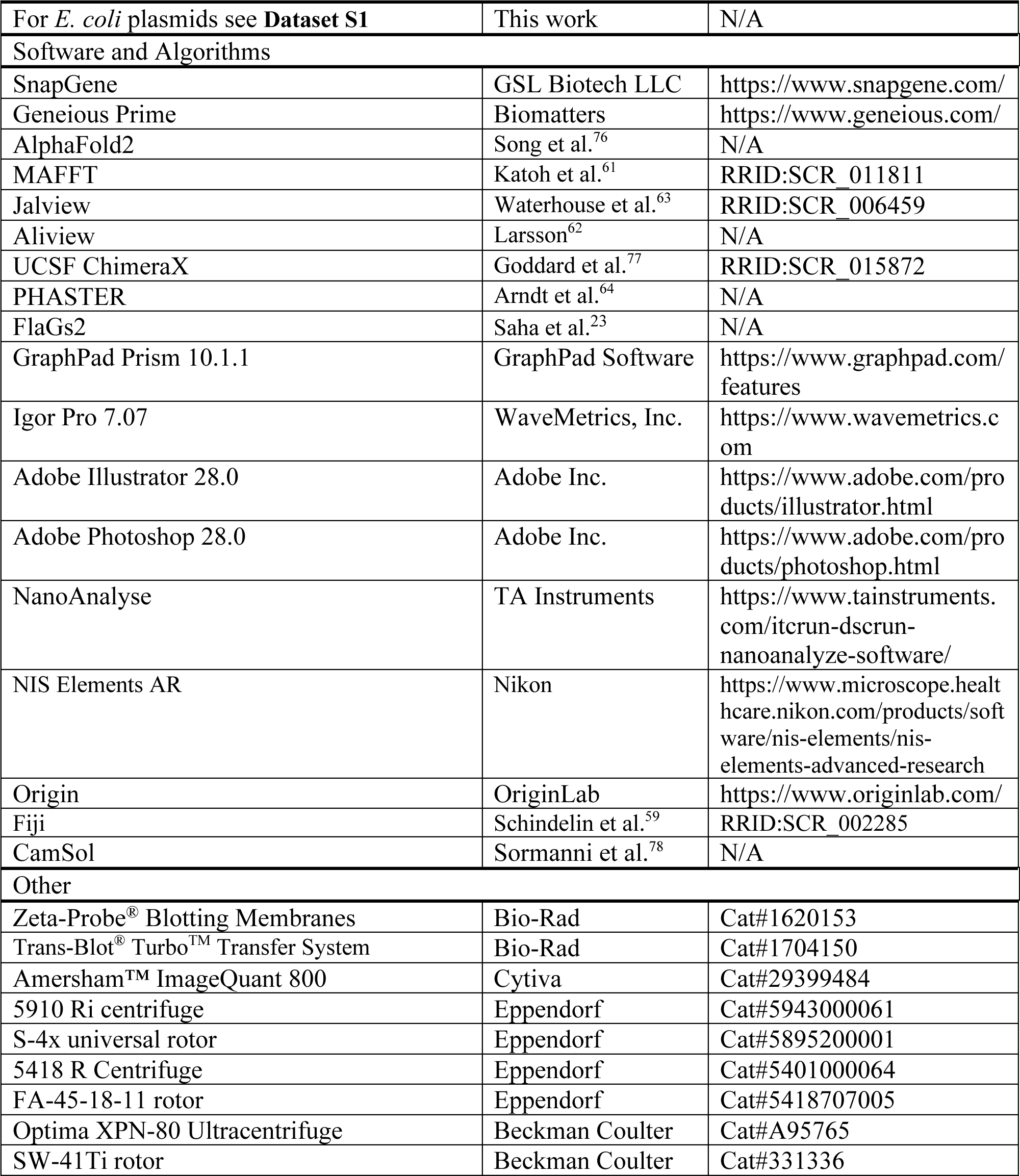

## SUPPLEMENTAL INFORMATION

Supplemental Information includes six figures and can be found with this article online at http://dx.doi.org/…

---

**Figure S1.**
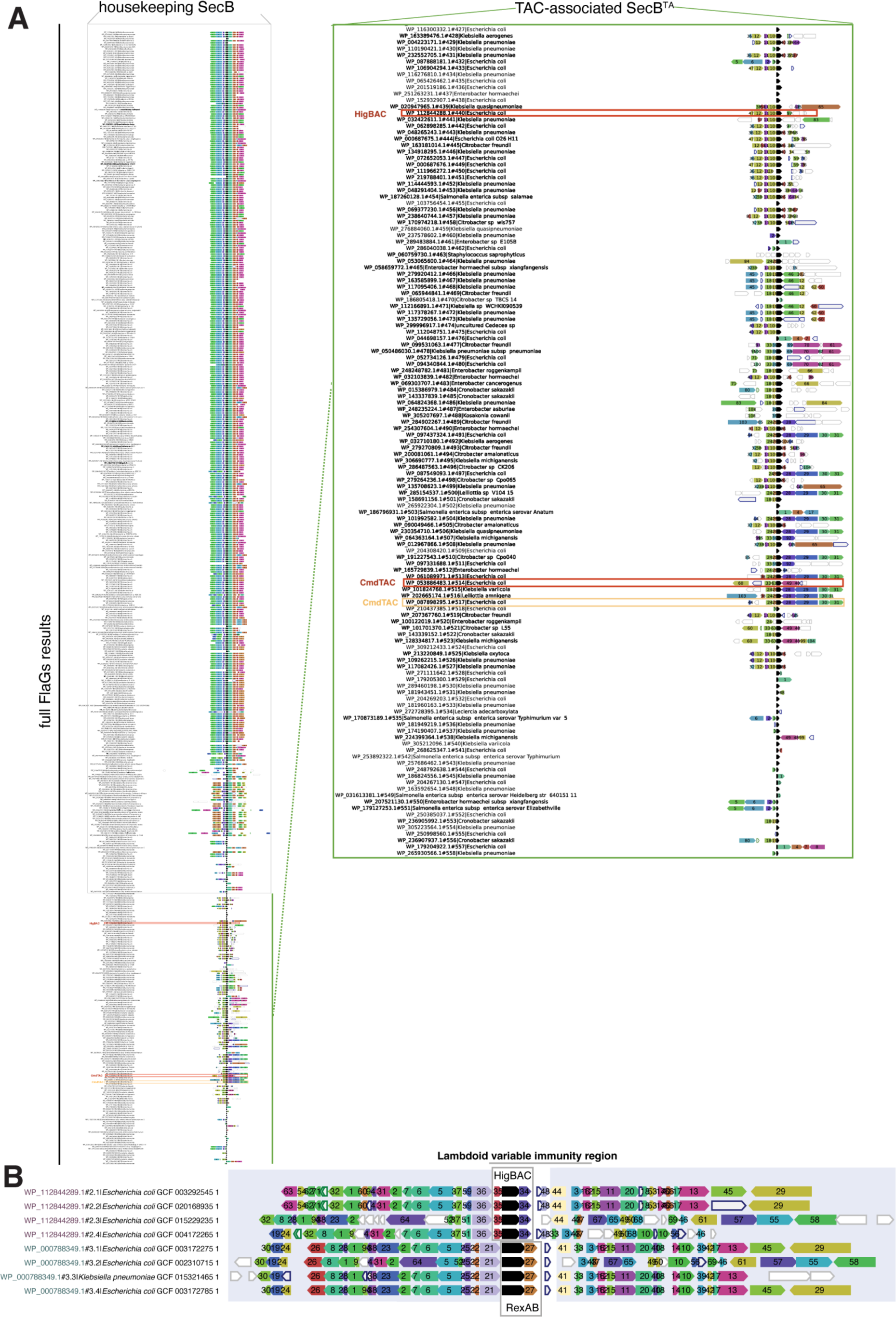
Genomic context of SecB homologues in Enterobacteriaceae. (**A**) Gene neighbourhood analysis of SecB genes in Enterobacteriaceae. Housekeeping SecB- like homologues are boxed in grey, while TAC-like SecBs are marked with green (zoomed in region in a green box). The TAC systems considered here are marked with red boxes, and the CmdTAC system described by Vassallo and colleagues^20^ is marked with an orange box (**B**) Gene neighbourhood analysis of the HigBAC system of Lambdoid prophages. Colours and numbers indicate genes encoding homologous proteins.

**Figure S2.**
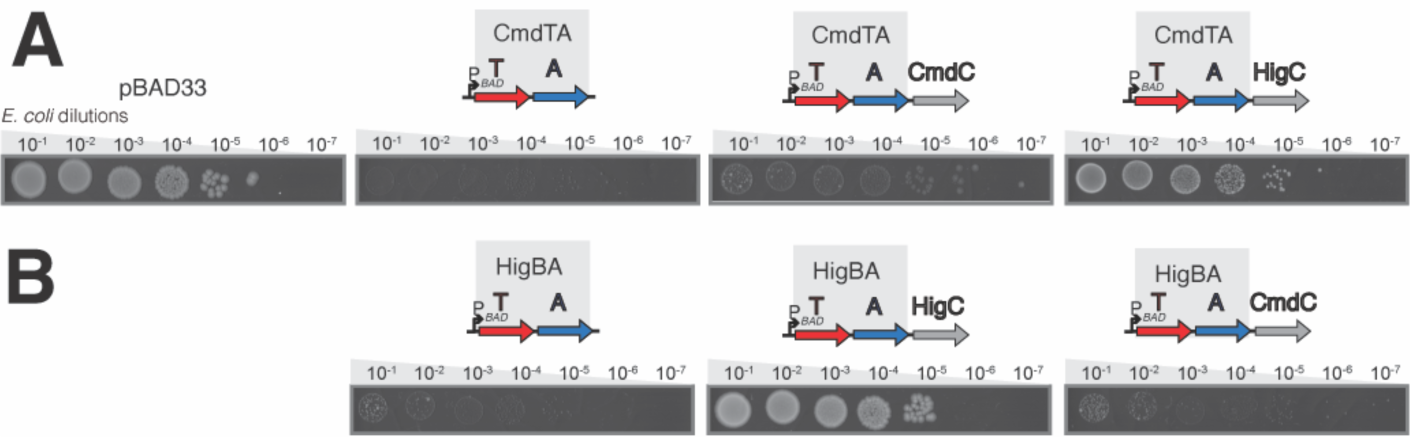
Neutralisation of TA units by cognate and non-cognate SecB^TA^ chaperones expressed as part of the TAC operon. *E. coli* BW25113 was transformed with either pBAD33 empty vector or pBAD33 derivatives expressing TA/TAC operons and grown for five to six hours. The bacterial culture was adjusted to OD600 1.0, serially diluted from 10^1^- to 10^8^-fold and spotted on LB medium supplemented with appropriate antibiotics and 0.2% arabinose for TA or TAC fragment induction.

**Figure S3.**
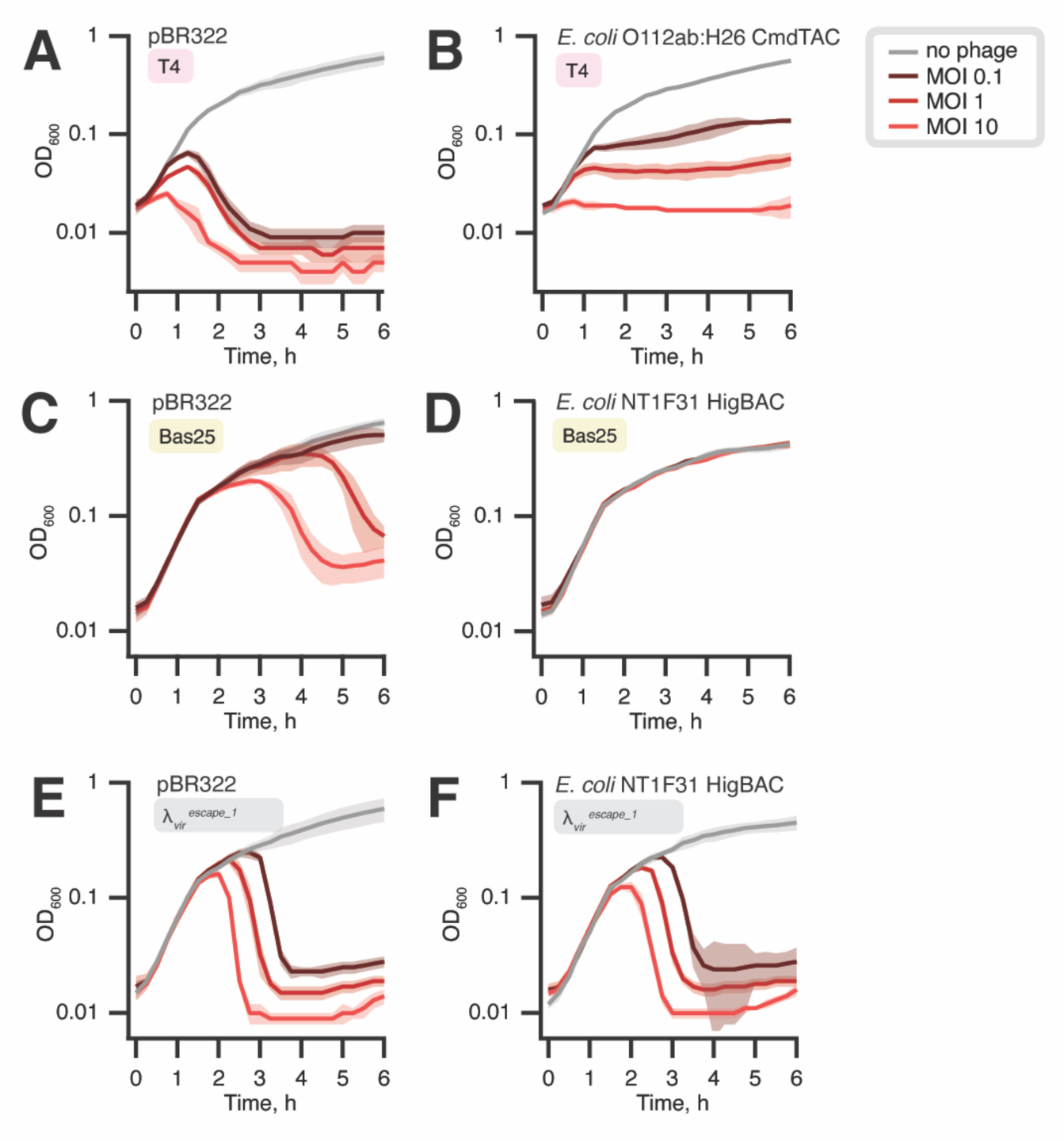
TAC systems display diverse modes of defence in liquid culture phage infection experiments. *E. coli* BW25113 transformed with either a pBR322 derivative lacking the tetracycline resistance cassette^30^ plasmid or a pBR322 derivative expressing a TAC defence system under the control of the P*tet* promoter were challenged with ten-fold serial dilutions of phages and grown at 37 °C in LB+Amp medium supplemented with 10 mM MgSO4 and 2.5 mM CaCl2. In the “no phage” experiments, the diluent was added instead of phages. Experiments were performed using three different transformants of each plasmid and the standard deviation is shown as a shadow. MOI stands for Multiplicity Of Infection, i.e. the ratio of phages to bacteria at the beginning of the experiment, time point 0.

**Figure S4.**
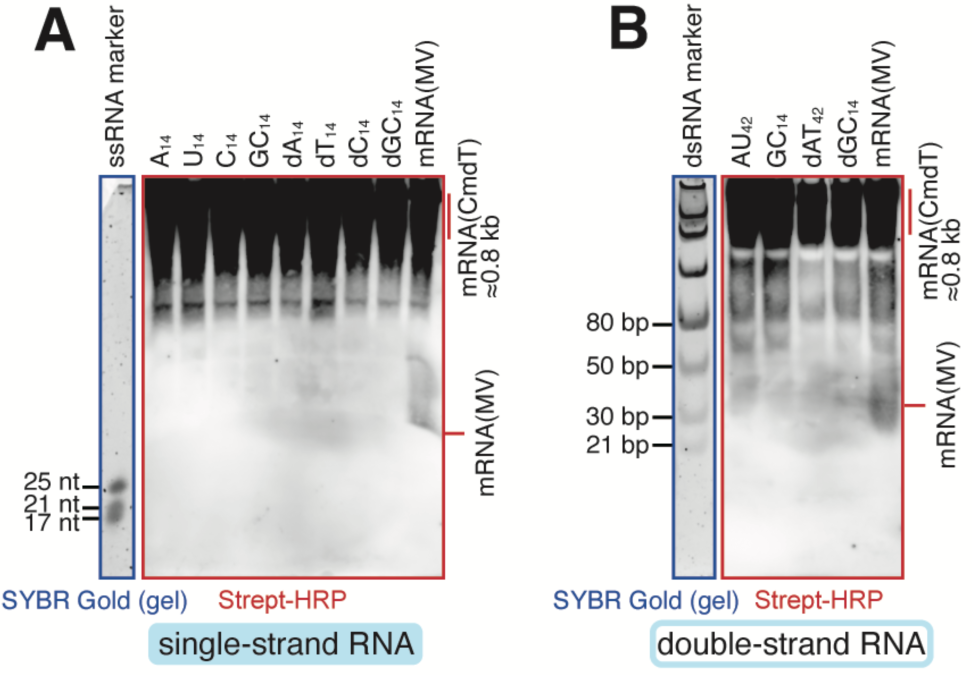
None of the tested homopolymeric RNA oligonucleotides are efficiently modified by *E. coli* O112ab:H26 CmdT in the presence of ^biot^NAD^+^. Experiments were performed using single-strand (**A**) and double-strand (**B**) RNA oligonucleotides. After the RNA modification reaction was completed, single-strand and double-strand RNA substrates were resolved in urea-PAGE and native PAGE, respectively. Samples prepared in the presence of ^biot^NAD^+^ were immunoblotted with streptavidin-HRP conjugate.

**Figure S5.**
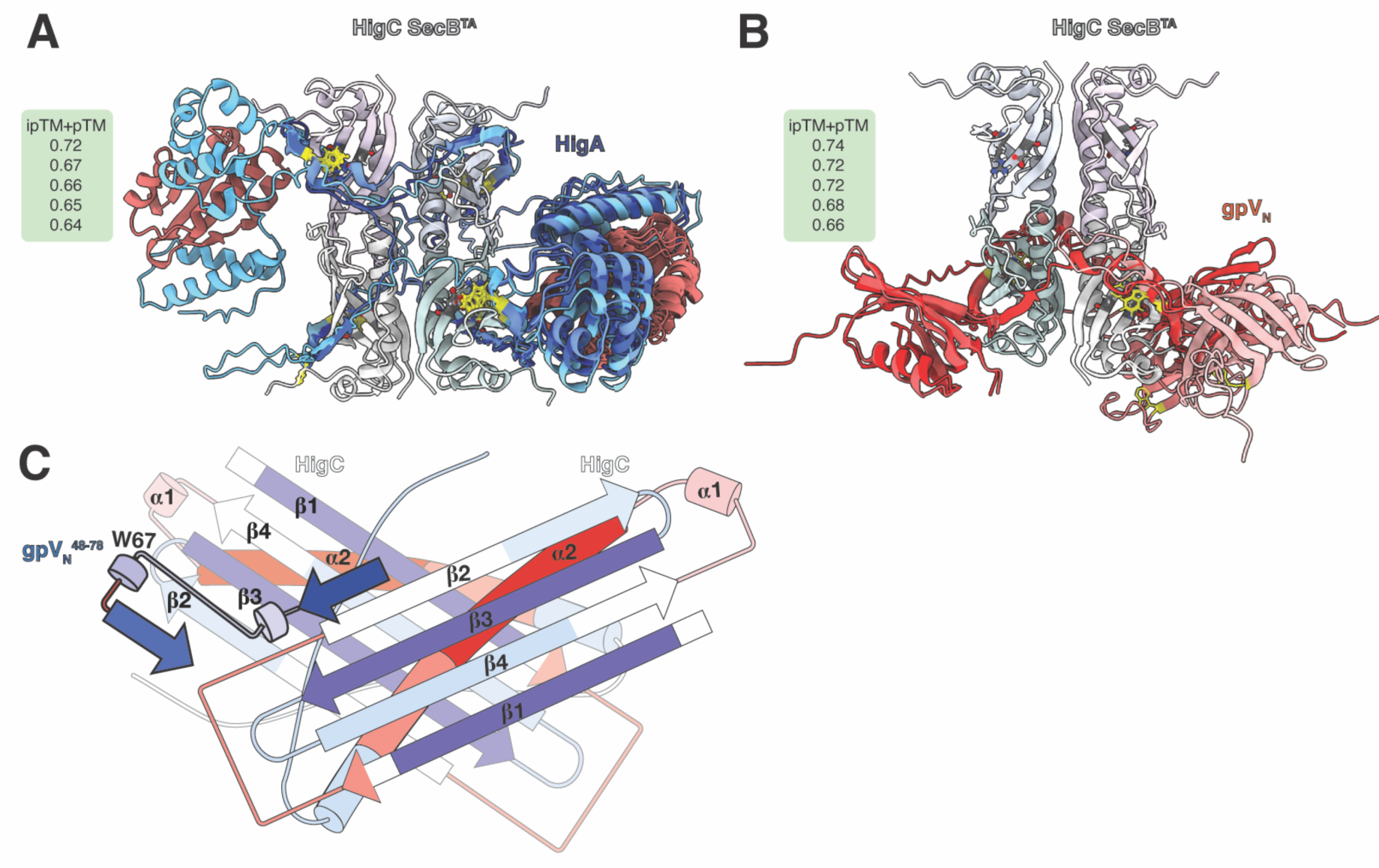
Structural modelling and ΔHDX analysis of HigC4 complexes. (**A,B**) Top-scoring AlphaFold models for SecB^TA^ complexes. Five top-scoring structural solutions for (**A**) HigA:HigB:HigC4 and (**B**) gpVN:HigC4. All structures are aligned using SecB^TA^. Only one chaperone tetramer structure is shown per structural alignment. pTM+ipTM scores are provided for the five top-scoring solutions. (**C**) Topology representation of HigC4:gpV^48–78^ colored as a function of the ΔHDX.

**Figure S6.**
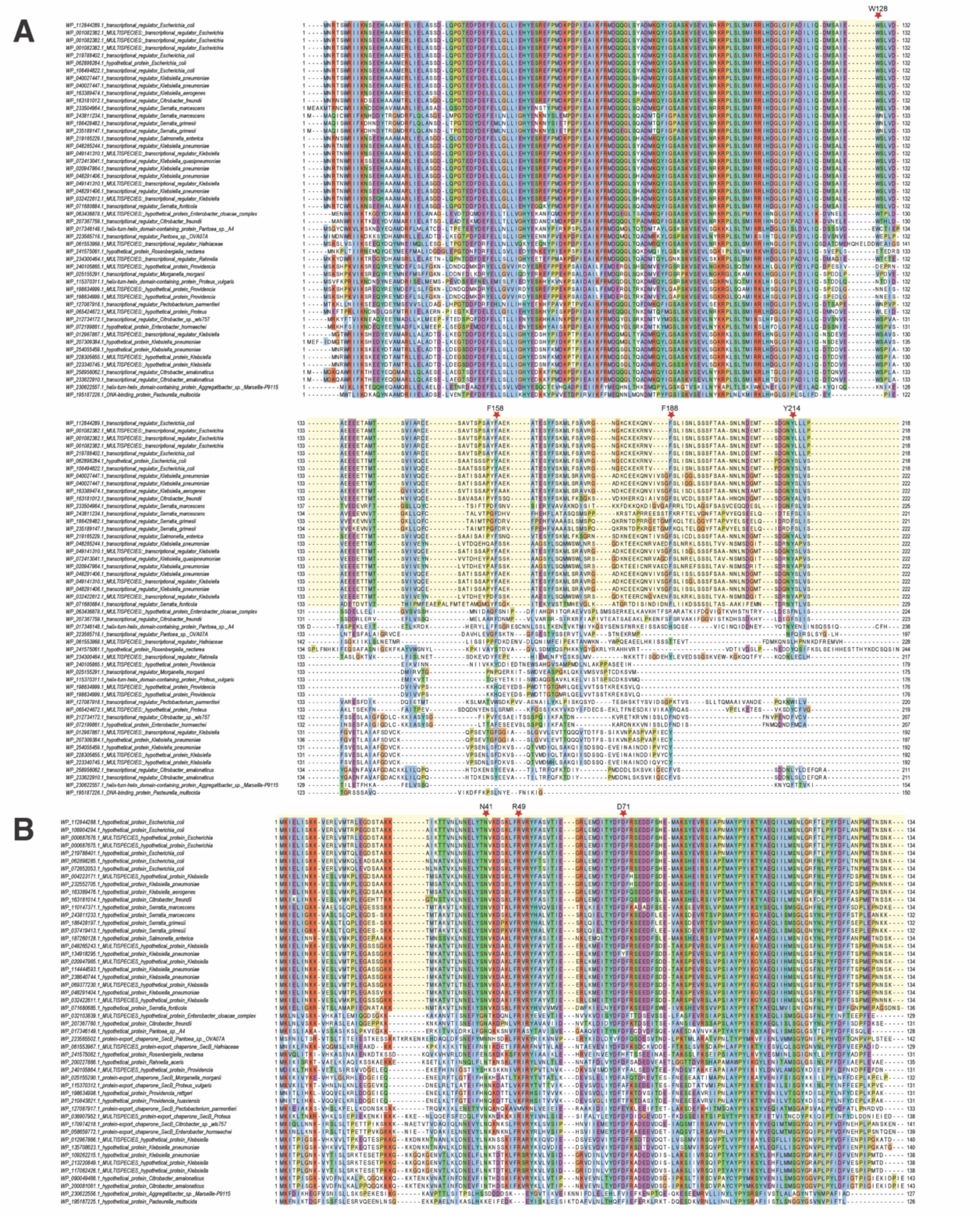
The ChAD is the most variable part of the HigA antitoxin, but is conserved in subgroups. (A) Alignment of HigBAC antitoxins. Red stars show the location of residue substitutions to alanine. Yellow block shading shows the subgroup with most similar ChADs. (**B**) Alignment of HigBAC chaperones. Red stars show key residues in the ChAD-chaperones interface. Most closely related SecB homologues to HigBAC chaperone and their respective HigBAC antitoxin ChADs are shaded in yellow.

## REFERENCES

1. 1. Jurėnas, D., Fraikin, N., Goormaghtigh, F., and Van Melderen, L. (2022). Biology and evolution of bacterial toxin-antitoxin systems. Nat Rev Microbiol 20, 335–350. 10.1038/s41579-021-00661-1.

2. LeRoux, M., and Laub, M.T. (2022). Toxin-Antitoxin Systems as Phage Defense Elements. Annu Rev Microbiol 76, 21–43. 10.1146/annurev-micro-020722-013730.

3. Song, S., and Wood, T.K. (2020). A Primary Physiological Role of Toxin/Antitoxin Systems Is Phage Inhibition. Front Microbiol 11, 1895. 10.3389/fmicb.2020.01895.

4. Kelly, A., Arrowsmith, T.J., Went, S.C., and Blower, T.R. (2023). Toxin-antitoxin systems as mediators of phage defence and the implications for abortive infection. Curr Opin Microbiol 73, 102293. 10.1016/j.mib.2023.102293.

5. Tian, Q.B., Ohnishi, M., Tabuchi, A., and Terawaki, Y. (1996). A new plasmid-encoded proteic killer gene system: cloning, sequencing, and analyzing hig locus of plasmid Rts1. Biochem Biophys Res Commun 220, 280–284. 10.1006/bbrc.1996.0396.

6. Schureck, M.A., Maehigashi, T., Miles, S.J., Marquez, J., Cho, S.E., Erdman, R., and Dunham, C.M. (2014). Structure of the *Proteus vulgaris* HigB-(HigA)2-HigB toxin-antitoxin complex. J Biol Chem 289, 1060–1070. 10.1074/jbc.M113.512095.

7. Christensen-Dalsgaard, M., and Gerdes, K. (2006). Two higBA loci in the Vibrio cholerae superintegron encode mRNA cleaving enzymes and can stabilize plasmids. Mol Microbiol 62, 397–411. 10.1111/j.1365-2958.2006.05385.x.

8. Hadži, S., Garcia-Pino, A., Haesaerts, S., Jurenas, D., Gerdes, K., Lah, J., and Loris, R. (2017). Ribosome-dependent Vibrio cholerae mRNAse HigB2 is regulated by a beta-strand sliding mechanism. Nucleic Acids Res 45, 4972–4983. 10.1093/nar/gkx138.

9. Jankevicius, G., Ariza, A., Ahel, M., and Ahel, I. (2016). The Toxin-Antitoxin System DarTG Catalyzes Reversible ADP-Ribosylation of DNA. Mol Cell 64, 1109–1116. 10.1016/j.molcel.2016.11.014.

10. LeRoux, M., Srikant, S., Teodoro, G.I.C., Zhang, T., Littlehale, M.L., Doron, S., Badiee, M., Leung, A.K.L., Sorek, R., and Laub, M.T. (2022). The DarTG toxin-antitoxin system provides phage defence by ADP-ribosylating viral DNA. Nat Microbiol. 10.1038/s41564-022-01153-5.

11. Suskiewicz, M.J., Prokhorova, E., Rack, J.G.M., and Ahel, I. (2023). ADP-ribosylation from molecular mechanisms to therapeutic implications. Cell 186, 4475–4495. 10.1016/j.cell.2023.08.030.

12. Schuller, M., Raggiaschi, R., Mikolcevic, P., Rack, J.G.M., Ariza, A., Zhang, Y., Ledermann, R., Tang, C., Mikoc, A., and Ahel, I. (2023). Molecular basis for the reversible ADP-ribosylation of guanosine bases. Mol Cell. 10.1016/j.molcel.2023.06.013.

13. Schuller, M., Butler, R.E., Ariza, A., Tromans-Coia, C., Jankevicius, G., Claridge, T.D.W., Kendall, S.L., Goh, S., Stewart, G.R., and Ahel, I. (2021). Molecular basis for DarT ADP-ribosylation of a DNA base. Nature 596, 597–602. 10.1038/s41586-021-03825-4.

14. Bordes, P., Cirinesi, A.M., Ummels, R., Sala, A., Sakr, S., Bitter, W., and Genevaux, P. (2011). SecB-like chaperone controls a toxin-antitoxin stress-responsive system in Mycobacterium tuberculosis. Proc Natl Acad Sci U S A 108, 8438–8443. 10.1073/pnas.1101189108.

15. Sala, A., Calderon, V., Bordes, P., and Genevaux, P. (2013). TAC from Mycobacterium tuberculosis: a paradigm for stress-responsive toxin-antitoxin systems controlled by SecB-like chaperones. Cell Stress Chaperones 18, 129–135. 10.1007/s12192-012-0396-5.

16. 16. Eismann, L., Fijalkowski, I., Galmozzi, C.V., Koubek, J., Tippmann, F., Van Damme, P., and Kramer, G. (2022). Selective ribosome profiling reveals a role for SecB in the co-translational inner membrane protein biogenesis. Cell Rep 41, 111776. 10.1016/j.celrep.2022.111776.

17. Texier, P., Bordes, P., Nagpal, J., Sala, A.J., Mansour, M., Cirinesi, A.M., Xu, X., Dougan, D.A., and Genevaux, P. (2021). ClpXP-mediated Degradation of the TAC Antitoxin is Neutralized by the SecB-like Chaperone in Mycobacterium tuberculosis. J Mol Biol 433, 166815. 10.1016/j.jmb.2021.166815.

18. Bordes, P., Sala, A.J., Ayala, S., Texier, P., Slama, N., Cirinesi, A.M., Guillet, V., Mourey, L., and Genevaux, P. (2016). Chaperone addiction of toxin-antitoxin systems. Nat Commun 7, 13339. 10.1038/ncomms13339.

19. Guillet, V., Bordes, P., Bon, C., Marcoux, J., Gervais, V., Sala, A.J., Dos Reis, S., Slama, N., Mares-Mejia, I., Cirinesi, A.M., et al. (2019). Structural insights into chaperone addiction of toxin-antitoxin systems. Nat Commun 10, 782. 10.1038/s41467-019-08747-4.

20. Vassallo, C.N., Doering, C.R., Littlehale, M.L., Teodoro, G.I.C., and Laub, M.T. (2022). A functional selection reveals previously undetected anti-phage defence systems in the *E. coli* pangenome. Nat Microbiol 7, 1568–1579. 10.1038/s41564-022-01219-4.

21. Ernits, K., Saha, C.K., Brodiazhenko, T., Chouhan, B., Shenoy, A., Buttress, J.A., Duque-Pedraza, J.J., Bojar, V., Nakamoto, J.A., Kurata, T., et al. (2023). The structural basis of hyperpromiscuity in a core combinatorial network of type II toxin-antitoxin and related phage defense systems. Proc Natl Acad Sci U S A 120, e2305393120. 10.1073/pnas.2305393120.

22. Maffei, E., Shaidullina, A., Burkolter, M., Heyer, Y., Estermann, F., Druelle, V., Sauer, P., Willi, L., Michaelis, S., Hilbi, H., et al. (2021). Systematic exploration of Escherichia coli phage-host interactions with the BASEL phage collection. PLoS Biol 19, e3001424. 10.1371/journal.pbio.3001424.

23. Saha, C.K., Sanches Pires, R., Brolin, H., Delannoy, M., and Atkinson, G.C. (2021). FlaGs and webFlaGs: discovering novel biology through the analysis of gene neighbourhood conservation. Bioinformatics 37, 1312–1314. 10.1093/bioinformatics/btaa788.

24. Salgado, H., Gama-Castro, S., Peralta-Gil, M., Diaz-Peredo, E., Sanchez-Solano, F., Santos-Zavaleta, A., Martinez-Flores, I., Jimenez-Jacinto, V., Bonavides-Martinez, C., Segura-Salazar, J., et al. (2006). RegulonDB (version 5.0): *Escherichia coli* K-12 transcriptional regulatory network, operon organization, and growth conditions. Nucleic Acids Res 34, D394–397. 10.1093/nar/gkj156.

25. Degnan, P.H., Michalowski, C.B., Babic, A.C., Cordes, M.H., and Little, J.W. (2007). Conservation and diversity in the immunity regions of wild phages with the immunity specificity of phage lambda. Mol Microbiol 64, 232–244. 10.1111/j.1365-2958.2007.05650.x.

26. Snyder, L. (1995). Phage-exclusion enzymes: a bonanza of biochemical and cell biology reagents? Mol Microbiol 15, 415–420. 10.1111/j.1365-2958.1995.tb02255.x.

27. Koubek, J., Schmitt, J., Galmozzi, C.V., and Kramer, G. (2021). Mechanisms of Cotranslational Protein Maturation in Bacteria. Front Mol Biosci 8, 689755. 10.3389/fmolb.2021.689755.

28. Wells, J.N., Bergendahl, L.T., and Marsh, J.A. (2016). Operon Gene Order Is Optimized for Ordered Protein Complex Assembly. Cell Rep 14, 679–685. 10.1016/j.celrep.2015.12.085.

29. Hurley, J.M., and Woychik, N.A. (2009). Bacterial toxin HigB associates with ribosomes and mediates translation-dependent mRNA cleavage at A-rich sites. J Biol Chem 284, 18605–18613. 10.1074/jbc.M109.008763.

30. Guegler, C.K., and Laub, M.T. (2021). Shutoff of host transcription triggers a toxin-antitoxin system to cleave phage RNA and abort infection. Mol Cell 81, 2361–2373 e2369. 10.1016/j.molcel.2021.03.027.

31. Kutter, E., Bryan, D., Ray, G., Brewster, E., Blasdel, B., and Guttman, B. (2018). From Host to Phage Metabolism: Hot Tales of Phage T4’s Takeover of E. coli. Viruses 10. 10.3390/v10070387.

32. Shimizu, Y., Kuruma, Y., Kanamori, T., and Ueda, T. (2014). The PURE system for protein production. Methods Mol Biol 1118, 275–284. 10.1007/978-1-62703-782-2_19.

33. Takada, H., Roghanian, M., Murina, V., Dzhygyr, I., Murayama, R., Akanuma, G., Atkinson, G.C., Garcia-Pino, A., and Hauryliuk, V. (2020). The C-terminal RRM/ACT domain is crucial for fine-tuning the activation of ‘long’ RelA-SpoT Homolog enzymes by ribosomal complexes. Frontiers in microbiology 11, 277. 10.3389/fmicb.2020.00277.

34. Evans, R., O’Neill, M., Pritzel, A., Antropova, N., Senior, A., Green, T., Žídek, A., Bates, R., Blackwell, S., Yim, J., et al. (2022). Protein complex prediction with AlphaFold-Multimer. bioRxiv, 2021.2010.2004.463034. 10.1101/2021.10.04.463034.

35. Jumper, J., Evans, R., Pritzel, A., Green, T., Figurnov, M., Ronneberger, O., Tunyasuvunakool, K., Bates, R., Zidek, A., Potapenko, A., et al. (2021). Highly accurate protein structure prediction with AlphaFold. Nature 596, 583–589. 10.1038/s41586-021-03819-2.

36. Xu, Z., Knafels, J.D., and Yoshino, K. (2000). Crystal structure of the bacterial protein export chaperone secB. Nat Struct Biol 7, 1172–1177. 10.1038/82040.

37. Holehouse, A.S., and Kragelund, B.B. (2023). The molecular basis for cellular function of intrinsically disordered protein regions. Nat Rev Mol Cell Biol. 10.1038/s41580-023-00673-0.

38. Sormanni, P., and Vendruscolo, M. (2019). Protein Solubility Predictions Using the CamSol Method in the Study of Protein Homeostasis. Cold Spring Harb Perspect Biol 11. 10.1101/cshperspect.a033845.

39. Flynn, J.M., Neher, S.B., Kim, Y.I., Sauer, R.T., and Baker, T.A. (2003). Proteomic discovery of cellular substrates of the ClpXP protease reveals five classes of ClpX-recognition signals. Mol Cell 11, 671–683. 10.1016/s1097-2765(03)00060-1.

40. Fei, X., Bell, T.A., Barkow, S.R., Baker, T.A., and Sauer, R.T. (2020). Structural basis of ClpXP recognition and unfolding of ssrA-tagged substrates. Elife 9. 10.7554/eLife.61496.

41. Shaner, N.C., Lambert, G.G., Chammas, A., Ni, Y., Cranfill, P.J., Baird, M.A., Sell, B.R., Allen, J.R., Day, R.N., Israelsson, M., et al. (2013). A bright monomeric green fluorescent protein derived from *Branchiostoma lanceolatum*. Nat Methods 10, 407–409. 10.1038/nmeth.2413.

42. Zhang, T., Tamman, H., Coppieters ’t Wallant, K., Kurata, T., LeRoux, M., Srikant, S., Brodiazhenko, T., Cepauskas, A., Talavera, A., Martens, C., et al. (2022). Direct activation of a bacterial innate immune system by a viral capsid protein. Nature 612, 132–140. 10.1038/s41586-022-05444-z.

43. Gao, L.A., Wilkinson, M.E., Strecker, J., Makarova, K.S., Macrae, R.K., Koonin, E.V., and Zhang, F. (2022). Prokaryotic innate immunity through pattern recognition of conserved viral proteins. Science 377, eabm4096. 10.1126/science.abm4096.

44. Campbell, P.L., Duda, R.L., Nassur, J., Conway, J.F., and Huet, A. (2020). Mobile Loops and Electrostatic Interactions Maintain the Flexible Tail Tube of Bacteriophage Lambda. J Mol Biol 432, 384–395. 10.1016/j.jmb.2019.10.031.

45. Pell, L.G., Kanelis, V., Donaldson, L.W., Howell, P.L., and Davidson, A.R. (2009). The phage lambda major tail protein structure reveals a common evolution for long-tailed phages and the type VI bacterial secretion system. Proc Natl Acad Sci U S A 106, 4160–4165. 10.1073/pnas.0900044106.

46. Liu, X., Jiang, H., Gu, Z., and Roberts, J.W. (2013). High-resolution view of bacteriophage lambda gene expression by ribosome profiling. Proc Natl Acad Sci U S A 110, 11928–11933. 10.1073/pnas.1309739110.

47. Xu, J., Hendrix, R.W., and Duda, R.L. (2014). Chaperone-protein interactions that mediate assembly of the bacteriophage lambda tail to the correct length. J Mol Biol 426, 1004–1018. 10.1016/j.jmb.2013.06.040.

48. Jurėnas, D., Payelleville, A., Roghanian, M., Turnbull, K.J., Givaudan, A., Brillard, J., Hauryliuk, V., and Cascales, E. (2021). *Photorhabdus* antibacterial Rhs polymorphic toxin inhibits translation through ADP-ribosylation of 23S ribosomal RNA. Nucleic Acids Res 49, 8384–8395. 10.1093/nar/gkab608.

49. Bullen, N.P., Sychantha, D., Thang, S.S., Culviner, P.H., Rudzite, M., Ahmad, S., Shah, V.S., Filloux, A., Prehna, G., and Whitney, J.C. (2022). An ADP-ribosyltransferase toxin kills bacterial cells by modifying structured non-coding RNAs. Mol Cell 82, 3484–3498 e3411. 10.1016/j.molcel.2022.08.015.

50. Stokar-Avihail, A., Fedorenko, T., Hor, J., Garb, J., Leavitt, A., Millman, A., Shulman, G., Wojtania, N., Melamed, S., Amitai, G., and Sorek, R. (2023). Discovery of phage determinants that confer sensitivity to bacterial immune systems. Cell 186, 1863–1876 e1816. 10.1016/j.cell.2023.02.029.

51. Schmitt, C.K., Kemp, P., and Molineux, I.J. (1991). Genes 1.2 and 10 of bacteriophages T3 and T7 determine the permeability lesions observed in infected cells of *Escherichia coli* expressing the F plasmid gene *pifA*. J Bacteriol 173, 6507–6514. 10.1128/jb.173.20.6507-6514.1991.

52. Labrie, S.J., Tremblay, D.M., Moisan, M., Villion, M., Magadan, A.H., Campanacci, V., Cambillau, C., and Moineau, S. (2012). Involvement of the major capsid protein and two early-expressed phage genes in the activity of the lactococcal abortive infection mechanism AbiT. Appl Environ Microbiol 78, 6890–6899. 10.1128/AEM.01755-12.

53. Huiting, E., Cao, X., Ren, J., Athukoralage, J.S., Luo, Z., Silas, S., An, N., Carion, H., Zhou, Y., Fraser, J.S., et al. (2023). Bacteriophages inhibit and evade cGAS-like immune function in bacteria. Cell 186, 864–876 e821. 10.1016/j.cell.2022.12.041.

54. Georgopoulos, C. (2006). Toothpicks, serendipity and the emergence of the Escherichia coli DnaK (Hsp70) and GroEL (Hsp60) chaperone machines. Genetics 174, 1699–1707. 10.1534/genetics.104.68262.

55. Zeilstra-Ryalls, J., Fayet, O., and Georgopoulos, C. (1991). The universally conserved GroE (Hsp60) chaperonins. Annu Rev Microbiol 45, 301–325. 10.1146/annurev.mi.45.100191.001505.

56. Xu, J., Hendrix, R.W., and Duda, R.L. (2004). Conserved translational frameshift in dsDNA bacteriophage tail assembly genes. Mol Cell 16, 11–21. 10.1016/j.molcel.2004.09.006.

57. 57. van der Vies, S.M., Gatenby, A.A., and Georgopoulos, C. (1994). Bacteriophage T4 encodes a co-chaperonin that can substitute for *Escherichia coli* GroES in protein folding. Nature 368, 654–656. 10.1038/368654a0.

58. Takano, T., and Kakefuda, T. (1972). Involvement of a bacterial factor in morphogenesis of bacteriophage capsid. Nat New Biol 239, 34–37. 10.1038/newbio239034a0.

59. Schindelin, J., Arganda-Carreras, I., Frise, E., Kaynig, V., Longair, M., Pietzsch, T., Preibisch, S., Rueden, C., Saalfeld, S., Schmid, B., et al. (2012). Fiji: an open-source platform for biological-image analysis. Nat Methods 9, 676–682. 10.1038/nmeth.2019.

60. Altschul, S.F., Madden, T.L., Schaffer, A.A., Zhang, J., Zhang, Z., Miller, W., and Lipman, D.J. (1997). Gapped BLAST and PSI-BLAST: a new generation of protein database search programs. Nucleic Acids Res 25, 3389–3402. 10.1093/nar/25.17.3389.

61. Katoh, K., and Standley, D.M. (2013). MAFFT multiple sequence alignment software version 7: improvements in performance and usability. Mol Biol Evol 30, 772–780. 10.1093/molbev/mst010.

62. Larsson, A. (2014). AliView: a fast and lightweight alignment viewer and editor for large datasets. Bioinformatics *30*, 3276–3278. 10.1093/bioinformatics/btu531.

63. Waterhouse, A.M., Procter, J.B., Martin, D.M., Clamp, M., and Barton, G.J. (2009). Jalview Version 2--a multiple sequence alignment editor and analysis workbench. Bioinformatics *25*, 1189-1191. 10.1093/bioinformatics/btp033.

64. Arndt, D., Grant, J.R., Marcu, A., Sajed, T., Pon, A., Liang, Y., and Wishart, D.S. (2016). PHASTER: a better, faster version of the PHAST phage search tool. Nucleic Acids Res 44, W16–21. 10.1093/nar/gkw387.

65. Zhang, Y., and Skolnick, J. (2004). Scoring function for automated assessment of protein structure template quality. Proteins 57, 702–710. 10.1002/prot.20264.

66. Hauser, M., Steinegger, M., and Soding, J. (2016). MMseqs software suite for fast and deep clustering and searching of large protein sequence sets. Bioinformatics 32, 1323–1330. 10.1093/bioinformatics/btw006.

67. Gu, Z. (2022). Complex heatmap visualization. iMeta 1, e43. doi.org/10.1002/imt2.43.

68. Schoch, C.L., Ciufo, S., Domrachev, M., Hotton, C.L., Kannan, S., Khovanskaya, R., Leipe, D., McVeigh, R., O’Neill, K., Robbertse, B., et al. (2020). NCBI Taxonomy: a comprehensive update on curation, resources and tools. Database (Oxford) 2020. 10.1093/database/baaa062.

69. Turner, D., Shkoporov, A.N., Lood, C., Millard, A.D., Dutilh, B.E., Alfenas-Zerbini, P., van Zyl, L.J., Aziz, R.K., Oksanen, H.M., Poranen, M.M., et al. (2023). Abolishment of morphology-based taxa and change to binomial species names: 2022 taxonomy update of the ICTV bacterial viruses subcommittee. Arch Virol 168, 74. 10.1007/s00705-022-05694-2.

70. Quan, J., and Tian, J. (2009). Circular polymerase extension cloning of complex gene libraries and pathways. PLoS One 4, e6441. 10.1371/journal.pone.0006441.

71. Quan, J., and Tian, J. (2011). Circular polymerase extension cloning for high-throughput cloning of complex and combinatorial DNA libraries. Nat Protoc 6, 242–251. 10.1038/nprot.2010.181.

72. Gibson, D.G., Young, L., Chuang, R.Y., Venter, J.C., Hutchison, C.A., 3rd, and Smith, H.O. (2009). Enzymatic assembly of DNA molecules up to several hundred kilobases. Nat Methods 6, 343–345. 10.1038/nmeth.1318.

73. Naville, M., Ghuillot-Gaudeffroy, A., Marchais, A., and Gautheret, D. (2011). ARNold: a web tool for the prediction of Rho-independent transcription terminators. RNA Biol 8, 11–13. 10.4161/rna.8.1.13346.

74. Neidhardt, F.C., Bloch, P.L., and Smith, D.F. (1974). Culture medium for enterobacteria. J Bacteriol 119, 736–747. 10.1128/jb.119.3.736-747.1974.

75. Grenier, F., Matteau, D., Baby, V., and Rodrigue, S. (2014). Complete Genome Sequence of Escherichia coli BW25113. Genome Announc 2. 10.1128/genomeA.01038-14.

76. Song, Y., DiMaio, F., Wang, R.Y., Kim, D., Miles, C., Brunette, T., Thompson, J., and Baker, D. (2013). High-resolution comparative modeling with RosettaCM. Structure 21, 1735–1742. 10.1016/j.str.2013.08.005.

77. Goddard, T.D., Huang, C.C., Meng, E.C., Pettersen, E.F., Couch, G.S., Morris, J.H., and Ferrin, T.E. (2018). UCSF ChimeraX: Meeting modern challenges in visualization and analysis. Protein Sci 27, 14–25. 10.1002/pro.3235.

78. Sormanni, P., Aprile, F.A., and Vendruscolo, M. (2015). The CamSol method of rational design of protein mutants with enhanced solubility. J Mol Biol 427, 478–490. 10.1016/j.jmb.2014.09.026.

